# Prevalent cross-cell type QTL *trans*-regulatory genetic effects impacting innate lymphoid cells in the small intestine

**DOI:** 10.1101/2025.04.16.649031

**Authors:** Miao Xu, Kirk Gosik, Yang Zang, Orr Ashenberg, Chenhao Li, Hongjun Li, Kai Liu, Gary Churchill, Daniel Gatti, Kwangbom Choi, Vivek Philip, Daniel Graham, Kushal K. Dey, Heping Xu, Ramnik J. Xavier, Aviv Regev

## Abstract

Tissue function in homeostasis and disease arise from coordinated interactions between different cell types and can vary between individuals in a population, in part due to the impact of genetic variants. In barrier tissues such as the gut, innate lymphoid cells (ILCs) play critical roles in maintaining tissue homeostasis and immunity, but the genetic basis and associated regulatory circuitry affecting tissue ILCs remain largely unknown. Here, we systematically mapped the regulatory functions of genetic variants across 273,370 gut ILCs profiled by single cell RNA-seq from 274 diversity outbred (DO) mice. Computational analysis identified quantitative trait loci (QTL) impacting ILCs at three levels: cell subtype-specific mRNA expression (local and distal), cell subset proportions, and ILC gene expression programs (in *trans*), of which we experimentally validated a pivotal role for the transcription factor *Rbpj* in regulating ILC3s. All three classes of ILC QTL display polygenic inheritance signatures and broadly overlap with QTL affecting peripheral cytokine levels that we measured in another 261 DO mice. Strikingly, nearly half of *trans*-QTL affecting ILC traits did not overlap (within a 50-kb window) genes expressed in ILCs, and such loci were enriched for genes expressed by non-ILCs especially enteric neurons and intestinal glia. This suggests that such loci may act instead across cell type boundaries and allowed us to recover causal *trans*-cell type regulatory circuits in the tissue, including loci encoding multiple neuron-expressed peptides, epithelial-cell expressed *Sox9*, and phagocyte-expressed *Ccl17*. Finally, human orthologs of genes in QTL impacting gene expression and cell proportions are enriched for autoimmune disease risk heritability, suggesting their relevance to human disease. Our findings highlight how the impact of genetic variants may propagate to maintain tissue homeostasis and show a path to understand causality in tissue biology using quantitative genetics.

## INTRODUCTION

The complex regulatory circuits within and across different cell types in a tissue impact cell–cell coordination, tissue homeostasis, function, and human disease^1^. While non-coding genetic variants initially contribute to variation in gene expression across individuals in a cell intrinsic manner, those changes likely propagate to affect tissue function through the impact of cells of one type on another^2^. However, the genetic basis of tissue function and homeostasis in complex mammalian organisms remains largely unknown, both in terms of overall principles and in specific mechanisms^3^.

A case in point is innate lymphoid cells (ILCs), early responders to tissue perturbation, which are critical for sustaining homeostasis at barrier surfaces, a process that is largely directed by cell subset-specific cytokine production^3–7^. The development and responses of ILCs are fine-tuned to adapt to different signals in barrier tissues, such as the intestines, to confer regulatory or effector function^8^ through subset-specific gene expression, changes in ILC composition, specialized gene programs^9–11^ and intercellular crosstalk^4^. Both physiological^4^ and genetic association^12^ studies have implicated ILCs in tissue dysfunction and human disease, including infectious, autoimmune, and neuronal diseases^13,14^. However, many of the genetic mechanisms underlying ILC circuits remain unknown.

Recent genetics and genomics approaches have been instrumental in deciphering gene function and circuits in both model systems and humans. Quantitative trait loci (QTL) studies, especially those applied to RNA^15–17^ (eQTL) or protein^18^ (pQTL) expression traits, have identified genetic variants and loci that impact the regulation of individual genes (in *cis* and *trans*) and gene programs (in *trans*)^1,2,18^. However, with notable recent exceptions^18,19^, most of these studies have been conducted with bulk profiling, confounding signals from multiple cell types in one tissue, which both reduces power and limits the ability to distinguish between cell intrinsic and extrinsic effects. Conversely, single cell genomics, especially single cell RNA-seq (scRNA-seq) has been used to chart atlases of gene expression in complex tissues at single cell resolution^20–22^, and has been combined with genome-wide association studies (GWAS) *post hoc*^23^ to relate disease genes to the cell types in which they are expressed or enriched^23–25^. However, such *post hoc* analysis has limited capacity for causal inference, and most scRNA-seq studies were either conducted at a scale that did not allow direct genetic associations, or were limited to non-tissue setting^19,26^, such as induced pluripotent stem cell (iPSC) lines or peripheral blood mononuclear cells (PBMCs).

scRNA-seq of tissue cells at the scale required for genetic association poses experimental and computational challenges and opportunities. Experimentally, there is a need to access and process a large number of samples efficiently. Computationally, methods are needed to perform genetic association not only for cell type specific gene expression, but also for gene programs and cell proportions, all recovered by scRNA-seq. Establishing such approaches can be facilitated by using meaningful biological model systems. A few recent studies have characterized immune phenotypes (but not single cell genomics profiles) in collaborative cross (CC) mice^27,28^. Another compelling system is diversity outbred (DO) mice, a heterogeneous stock derived from outbreeding mice from eight founder strains (C57BL/6J, 129S/SvlmJ, PWK/PhJ, NOD/ShiLtJ, A/J, NZO/HlLtJ, CAST/ EiJ, WSB/EiJ)^29,30^ with available whole genome sequence data. This mouse population presents balanced allele frequencies, simple population structure, and extensive natural genetic variants within a relatively small sample size required for mapping compared to human populations^30^. Importantly, it captures genetic diversity and predisposition at both the cell and organism levels and can serve as a more compelling model than standard laboratory inbred strains^31^.

Here, we performed multiplexed scRNA-seq of 273,370 ILCs from the small intestine of 274 outbred, genotyped DO mice, along with cytokine profiling in another 261 DO mice, and developed a computational framework to identify QTL impacting ILC subtype specific gene expression (locally/in *cis* and distally/in *trans*), gene programs (in *trans*) and proportions (in *trans*). Hundreds of novel QTL were associated with ILC traits in a polygenic architecture, 34 of which also impacted peripheral cytokine levels. Of those associations, we experimentally validated *in vivo* the role of *Rbpj* in regulating intestinal ILC3 proportions and cytokine production. Remarkably, nearly half (48%) of the QTL affecting ILC traits in trans did not overlap (within a 50-kb window) any genes expressed in ILCs, and instead include genes expressed in other intestinal cell types, especially enteric neurons and intestinal glia. These suggest multiple causal cell–cell interactions from neural, immune and epithelial cells to ILCs, highlighting ways by which the impact of a genetic variant is causally propagated in tissue. Thus, our work provides a scalable approach to decipher the impact of genetic variants on tissue function.

## RESULTS

### Multiplexed scRNA-seq of intestinal ILCs from DO mice

Using multiplexed scRNA-seq, we collected 273,370 high-quality ILC profiles from the small intestine of 274 DO mice. We isolated total intestinal ILCs from DO mice by fluorescence-activated cell sorting (FACS) and performed scRNA-seq in multiplexed pools (**Fig. 1A, Methods**), which we subsequently demultiplexed^32^ based on genetic variants (**Methods**). In parallel, we genotyped each DO mouse with a GIGA-MUGA genotyping microarray^33^, capturing 143,179 SNPs across the mouse genome (**Fig. S1A**). We integrated the data from different experimental batches, followed by unsupervised clustering and *post hoc* annotation by known signature genes (**Fig. S1B**). In particular, among 371,654 profiled cells we annotated 273,370 cells as canonical ILC subsets and states: ILC1s, ILC2s, ILC3s (RORγt^high^/RORγt^low^) and ILC3-like LTis (CCR6^high^/CCR6^low^), hereafter referred to as “LTi”^5,34^ (**Fig. 1B, C**). Other cell types were captured at smaller proportions in our sort, including DCs, B cells, NKT cells and enterocytes (**Fig. 1B**) and filtered from subsequent analyses, unless specified.

**Fig. 1.**
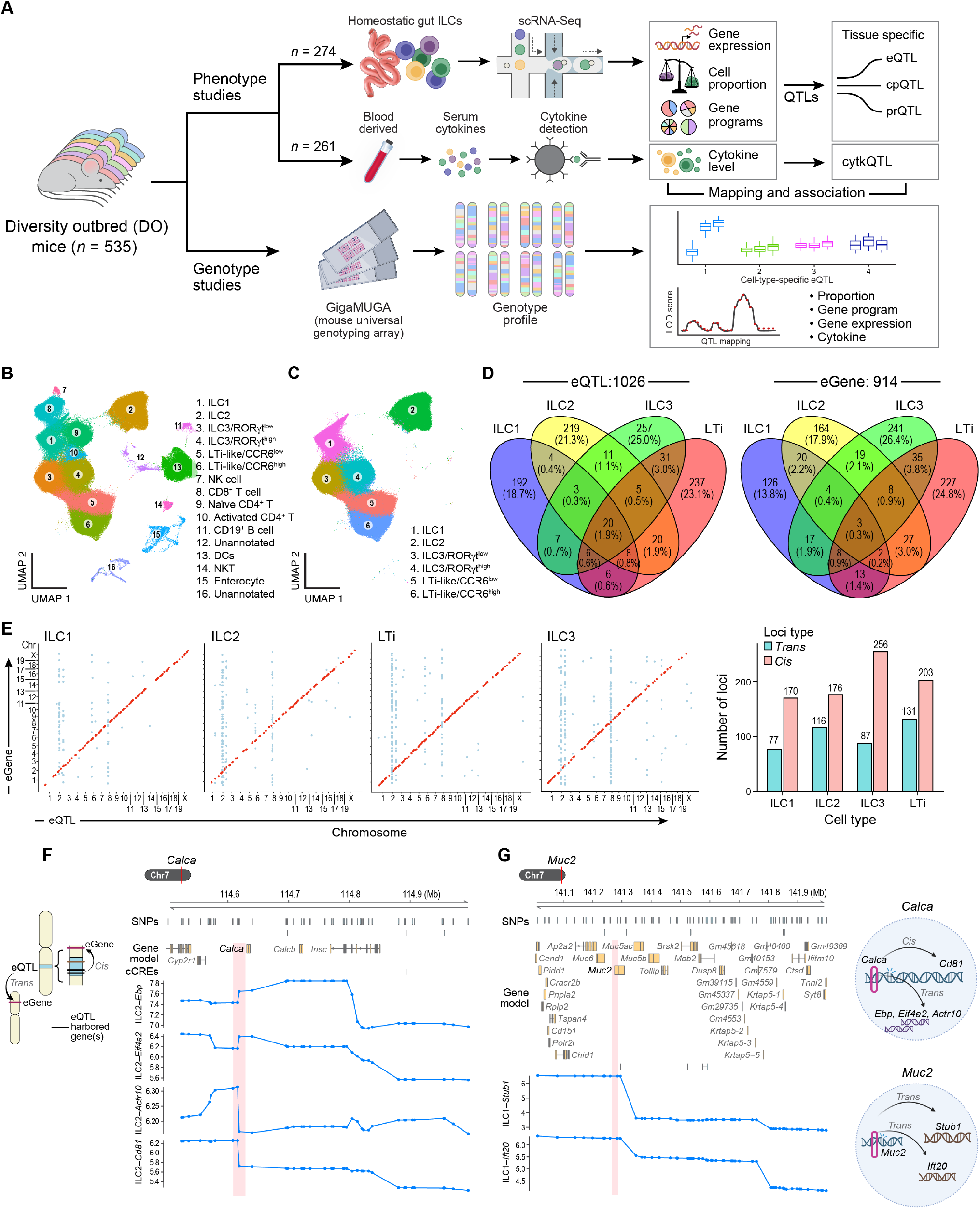
Multiplexed scRNA-seq in DO mice identifies shared and specific eQTLs in ILC subsets. (**A**) Study overview. (**B, C**) ILC cell atlas. Uniform Manifold Approximation and Projection (UMAP) embedding of cell profiles (dots) of cells isolated from the small intestine and enriched with FACS for ILCs **(B)** or only those annotated as ILCs (**C**), colored by clustering, and annotated *post hoc*. (**D**) ILC subset specific and shared eQTLs and eGenes. Venn diagram of eQTLs (left) and the eGenes whose expression they affect (right) from analysis in each ILC subset. (**E**) *Cis*- and *trans*-eQTL. Left: eQTL associations (dots) between a genetic variant (eQTL, *x*-axis) and affected eGene (*y*-axis) for each ILC subset (panels). Red: *cis*-eQTLs (diagonal); blue: *trans*-eQTLs (eGenes > 1 Mb from eQTL or on a different chromosome; off diagonal). Right: Number of *cis*-(red) and *trans-* (blue) eQTLs in each ILC subset. (**F, G**) Example eQTLs. From top: Gene models (top), conserved regulatory elements (CRE) and LOD scores (*y*-axis) of QTL effects in a locus including *Calca* (red vertical stripe: eQTL) associated with variation in ILC2 expression of *Ebp, Eif4a2* and *Actr10* in *trans* and *Cd81* in *cis* (**F**) and a locus including *Muc2* (red vertical stripe: eQTL) associated with ILC1-expression of *Stub1* and *Ift20* (in *trans*) (**G**). Right: illustrative model of *cis* and *trans* effects.

### Shared and specific eQTL in ILC subsets

eQTL mapping (**Methods**) in each of the four ILC subsets separately identified 1,026 eQTL (genome-wide significance *p* < 0.05) affecting the expression of 914 ILC genes, termed eGenes^35^ (**Fig. 1D**). Most eQTL (88.2%) and eGenes (82.9%) were ILC subset-specific, while the rest were shared by two or more ILC subsets (**Fig. 1D**). Most (66.2%) eQTL were local (“*cis*-eQTL”) associated with eGene traits within 1 Mb or less (**Fig. 1E**, red), and the remainder acted distally^36–38^, often on multiple genes across different chromosomes (**Fig. 1E**, blue) (“*trans*-eQTL”), with 4.3% acting both locally and distally. In particular, *trans*-associated loci on chromosome 2 and 8 affected many eGenes across multiple chromosomes in multiple subsets (**Fig. 1E**). Most eQTL spanned enhancer regions as annotated by ENCODE^39^, suggesting a regulatory effect on expression^40,41^ (**Fig. S1C, Methods**).

We further analyzed the genes overlapping ILC eQTL (local and distal), which we termed eQTL-overlapping genes (**Methods**). For example, *Calca*, encoding calcitonin gene-related peptide (CGRP) and expressed in ILC2s (**Fig. 1D**), overlaps an eQTL impacting eGenes *Cd81* (locally), *Ebp, Eif4a2*, and *Actr10* (in *trans*) in ILC2s (**Fig. 1F**). CGRP is a neuropeptide that plays a key role in type II immunity and intestinal homeostasis through restraining ILC2 expansion and promoting IL-5 expression^10,13,42^. Excess activation of CD81^+^ ILC2s leads to inflammation under chronic cold stress^45^. In another example, *Muc2*, encoding MUC2, overlaps a *trans*-eQTL locus in chromosome 7 that affects the expression of eGenes *Stub1* (on Chr 17) and *Ift20* (on Chr 11) in ILC1s (**Fig. 1G**). MUC2, expressed by enterocytes^46^ (**Fig. S1E**), is the predominant gel-forming mucin that contributes to the formation of the protective mucus barrier^47^. eGene *Stub1* is an E3 ligase implicated in gastric cancer^48^, and *Ift20* encodes a transport protein that recruits ATG16L1 to promote autophagosome biogenesis and is a key susceptibility gene for inflammatory bowel disease (IBD)^49,50^.

Overall, we identified a substantial number of *cis*- and *trans*-associations between ILC subset specific and shared eQTL and gene expression.

### Variants in loci for key ILC and tissue homeostasis regulators, including *Fam21, Il20ra1* and *Rpbj*, affect the balance of ILC subsets

Imbalances in the proportions of different ILC subsets in either local tissue or peripheral circulation have been associated with pathogenic immune responses, such as inflammatory arthritis^51^, septic shock^52^ and lupus^53^, as well as disease susceptibility and severity^54–56^ in chronic obstructive pulmonary disease (COPD)^57^ and allergy^58^.

To investigate the genetic basis of ILC subset proportions, we first estimated the composition of ILC subsets by scRNA-seq, finding substantial variation across individual DO mice (**Fig. 2A, Fig. S2A**), and then tested for loci associated with ILC sub-set ratios. We found 173 cell proportion QTL (cpQTL, *P* < 10^−6^, genome-wide significance, **Methods**), defined as loci that af-fect the ratio between at least one pair of ILC subsets: ILC1/ILC2, ILC1/ILC3, ILC2/ILC3, ILC2/LTi, ILC3/LTi and ILC1/LTi (**Fig. 2B, Fig. S2B**). (Note that these traits are not neces-sarily independent.) Additionally, 68 QTL were associated with the relative proportion of functionally distinct^59^ subsets within ILC3s (RORγt^high^ vs. RORγt^low^) or LTis (CCR6^high^ vs. CCR6^low^) (**Fig. 2C**).

**Fig. 2.**
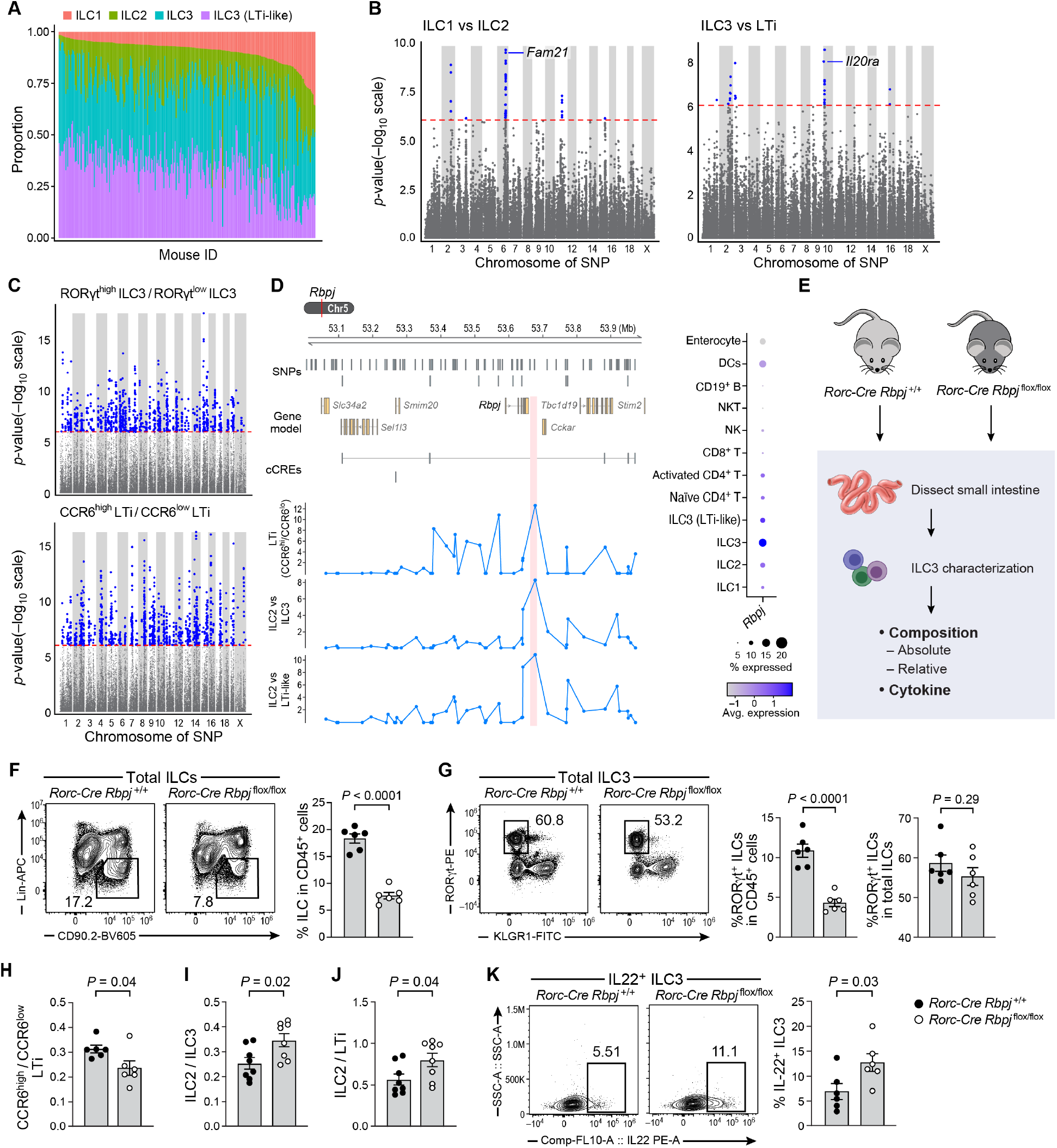
Proportion QTLs affecting the balance of ILC subsets overlap key ILC and tissue homeostasis regulators. (**A**) ILC subset proportions vary across DO mice. Proportion (*y*-axis) or each major ILC subset (color) across DO mice (*x*-axis), sorted by ILC1 proportion. (**B**,**C**) Proportion QTLs. Significance (*y*-axis, –log_10_ (*P*-value)) of association of each genomic locus (*x*-axis) with ratio of ILC1 vs. ILC2 (**B**, left), ILC3 vs. LTi (**B**, right), RORγt^high^ILC3 vs. RORγt^low^ ILC3 (**C**, top) and CCR6^high^ LTi vs. CCR6^low^ LTi (**C**, bottom). Dotted red line: genome-wide significance *p*-value = 0.05 (Bonferroni corrected). Blue: significant associations. (**D**) Proportion QTLs in the *Rpbj* locus. Left: Gene models (top), conserved regulatory elements (CRE) and LOD scores (*y*-axis) of QTL effects in a locus including *Rpbj* (red vertical stripe: proportion QTL) and associated with variation in proportion of CCR6^high^/CCR6^low^ LTis, ILC2/ILC3, and ILC2/LTi. Right: Mean expression (dot color; log(TPM+1)) and proportion of expressing cells (dot size) for *Rbpj* across different cell subsets (rows) in DO mice. (**E–K**) Validation of the role of *Rbpj* in ILC3 regulation in homeostasis through *Rbpj* conditional knockout mice. (**E**) Experimental approach. (**F**) Reduced total ILCs in *Rpbj* cKO mice. Left: Proportion of total ILCs (gated box) in CD45^+^ cells from the small intestine of wild-type *Rbpj*^+/+^*Rorc*^Cre^ (left-most) and *Rbpj*^flox/flox^*Rorc*^Cre^ (second from left) mice. Right: Proportion of total ILCs in CD45^+^ cells (*y*-axis) in wildtype (black circle) and cKO (white circles) mice (*x*-axis). *n* = 6 individual mice in each category (dots). (**G**) Reduced ILC3s (RORγt^+^ ILC) among total CD45^+^ cells and ILCs in *Rpbj* cKO mice. Proportion of RORγt^+^ ILC (gated box) in total CD45^+^ cells from the small intestine of wild-type *Rbpj*^+/+^*Rorc*^Cre^ (left-most) and *Rbpj*^flox/flox^*Rorc*^Cre^ (second from left) mice. Right: Proportion of RORγt^+^ ILC from total ILCs (*y*-axis second from right) or from CD45^+^ cells (right, *y*-axis) in wildtype (black circles) and cKO (white circles) mice (*x*-axis). *n* = 6 individual mice in each category (dots). (**H–J**) Changes in other ILC subset proportions in *Rpbj* cKO mice. Relative proportion of CCR6^+^/CCR6^−^ LTi (*y*-axis, **H**), ILC2/ILC3 (*y*-axis, **I**), and ILC2/LTi (*y*-axis, **J**) in wildtype (black circles) and cKO (white circles) mice (*x*-axis). *n* = 6 individual mice in each category (dots). (**K**) Increase in IL-22 producing ILC3s in *Rpbj* cKO mice. Proportion of IL22^+^ ILC3s (gated box) in total ILC3s from the small intestine of wild-type *Rbpj*^+/+^*Rorc*^Cre^ (left-most) and *Rbpj*^flox/flox^*Rorc*^Cre^ (second from left) mice. Right: Proportion of IL22^+^ ILC3s from total ILC3s (*y*-axis) in wildtype (black circles) and cKO (white circles) mice (*x*-axis). *n* = 6 individual mice in each category (dots). Flow cytometry panels (**F–K)** are representative of at least two independent experiments. Error bars: SEM. *P*-value: unpaired two-tailed *t* test.

Some of the putative causal genes in the associated cpQTL regions are implicated in key processes relevant to ILC regulation and tissue homeostasis. For example, *Fam21* (*Washc2*), which overlaps one of 17 loci associated with the ILC1/ILC2 ratio (**Fig. 2B, Fig. S2C**), encodes a known regulator of the endosomal localization of the WASH regulatory complex, which is in turn essential to sustain the intestinal NKp46^+^ ILC3 pool at homeostasis^60^. Our finding suggests an additional role in this context. Notably, *Fam21* is most highly expressed in DCs (where it is essential for the defense from *Candida albicans*^61^) and lymphocytes, but very lowly expressed in ILCs (**Fig. S2C**). In another example, *Il20ra* is proximal (along with other genes) to one of the 38 QTL associated with the ILC3 vs. LTi ratio (**Fig. 2B, Fig. S2D**). The ILC3 vs. LTi ratio accounts for differ-ential IL17 (pro-inflammatory) to IL-22 (protective) immune responses, respectively^62^. The human IL20ra locus (6q23) is strongly associated with autoimmune diseases, and *Il20ra* is its causal gene, as demonstrated by long-range chromatin interac-tion in T cells^63,64^. Our analysis suggests that in mouse *Il20ra* is both associated with the ratio of ILC3 vs. LTi and is highly expressed in these cells (as well as in T cells and ILC1s) (**Fig. S2D**).

Notably, the TF *Rbpj* is proximal (with other genes) to a QTL significantly associated with variations in several subset ratios, including ILC2/ILC3, ILC2/LTi and CCR6^high^LTi/CCR6^low^ LTi (**Fig. 2D**). *Rbpj* is a DNA-binding protein that associates with all Notch isoforms and mediates the transcriptional output of Notch signaling. Notch signaling is required for development of adult RORγt^+^ ILCs, including intestinal NKp46+ ILCs (ILC3) and some LTi-like ILCs^65–68^. In our data, Rpbj is expressed in a higher fraction of ILC3s and LTis vs. other ILCs and all its associated trait proportions involve ILC3s and/or LTis (**Fig. 2D**). We thus reasoned that *Rpbj* might play a role in specifically regulating ILC3s.

To test this hypothesis, we generated *Rorc*^Cre^*Rbpj*^fl/fl^ conditional knock-out mice (*Rbpj*^cKO^), where *Rbpj* is preferentially depleted in total *Rorc*-expressing ILC3s (conventional ILC3s and ILC3-like LTis), and compared their cell composition and cytokine secretion to those of *Rorc*^Cre^*Rbpj*^+/+^ mice (*Rbpj*^WT^) (**Fig. 2E, Methods**). We isolated total small intestinal CD45^+^ cells and assessed the composition of ILC subsets by defined markers as: total ILCs (CD45^+^Lin^−^CD90.2^+^), total ILC3s (RORγt^+^ ILCs: conventional ILC3s and LTis), convention-al ILC3s (RORγt^+^NKP46^+^CCR6^−^ ILCs), and ILC3-like Ltis (RORγt^+^NKP46^−^CCR6^+^ ILCs) (Methods). *Rbpj*^cKO^ mice had significantly lower proportions of total ILCs, total ILC3s and conventional ILC3s out of all CD45^+^ cells (**Fig. 2F, G, Fig. S2E**), lower proportions of total and conventional ILC3s out of all ILCs, and lower proportions of conventional ILC3 out of all ILC3s (**Fig. 2G, Fig. S2E, F**). Notably, the three proportion changes in our DO mice trait analysis — CCR6^high^LTi/CCR6^low^ LTi, ILC2/ILC3 and ILC2/LTi (RORγt ^+^NKP46^−^) — were sig-nificantly impacted in the *Rbpj*^cKO^ mice (**Fig. 2D, H–J**). (Note however that in the cpQTL all three proportions are impacted in the same (positive) direction with the minor allele, but in the KO setting, the proportion CCR6^high^ LTi/CCR6^low^ LTi shifted in the opposite direction than the other two.) To determine the im-pact of *Rbpj* on local cytokine production, we measured intra-cellular cytokine abundance and found that *Rbpj*^cKO^ ILC3s had significantly higher IL-17A and IL-22 levels compared to those from *Rbpj*^WT^ mice (**Fig. 2K, Fig. S2G, Methods**). Thus, RBPJ, a cell proportion QTL gene, regulates both intestinal ILC3 com-position and their cytokine production.

### Genes encoding regulators of intestinal function overlap QTL impacting ILC gene programs

Gene programs consisting of multiple co-regulated genes^69^ can characterize cell states, subsets or shared processes^9,10^ and be associated with context-dependent *trans* QTL^70^. To test for genetic variation associated with differences in gene program expression, we first used topic modeling to define 20 gene programs^70,66^ (**Fig. 3A, Methods**) that are either active in one ILC subset or across cells from multiple subsets (**Fig. 3B, Fig. S3A**). For example, topic 11 (“ILC2s”, **Fig. 3B**), scored highly in ILC2s expressing its signature TFs *Gata3* and *Calca* (**Fig. 3C**). Topic 0 (“ILC1s”, Fig. 3B) had 85 high scoring genes (e.g., *Ifngr1, Ncr1, Junb, Il2rb, Ccl5, Ccl3*, and *Ccl4*), was highly expressed in ILC1s, and was enriched for gene sets involved in ILC1 functions, such as response to IFNγ and regulation of cytotoxicity^71^ (**Fig. 3D**). Topics 3, 8, 12 and 16 each scored highly in a different portion of the ILC3s and/or LTis phenotypic space (e.g., Topic 3 in CCR6^high^ LTis; topic 16 in CCR6^low^ LTis) and were enriched for genes sets involved in both shared processes (myeloid cell differentiation and viral gene expression) and in distinct ones (intrinsic apoptotic pathways and response to IL-4 or hyperoxia in CCR6^low^ LTi topic 16; myeloid cell homeostasis, regulation of NF-κB signaling, response to TGF-β in CCR6^high^ LTi topic 3; **Fig. 3B, D**).

**Fig. 3.**
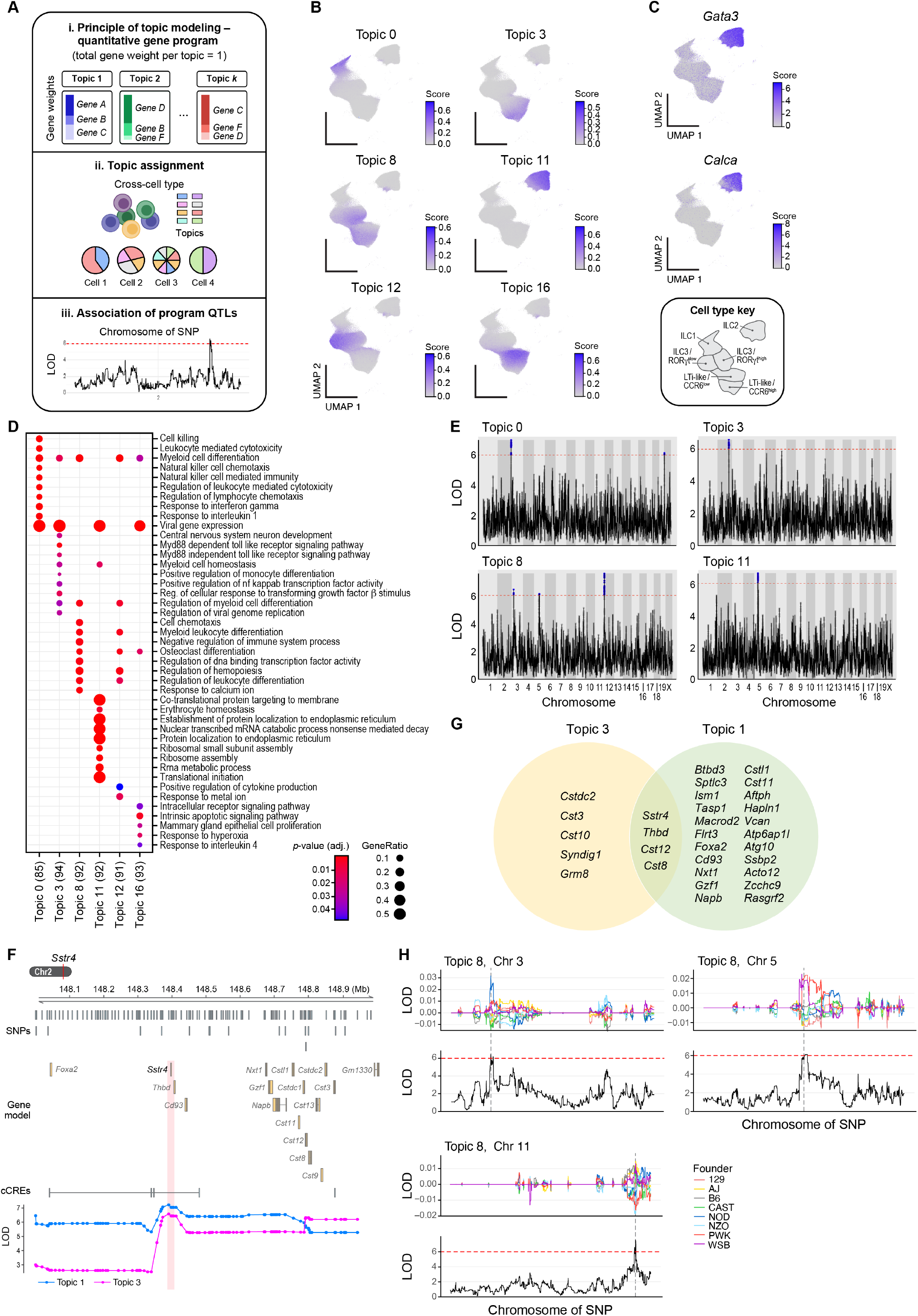
Program QTLs overlap regulators of intestinal function. (**A**) Topic models and program QTL approach overview. (**B–D**) Cell type specific programs. (**B**) UMAP embedding of ILC profiles (as in **Fig. 1C**), colored by cell scores for topics 0, 3, 8, 11, 12 and 16, or by expression of *Gata3* (encoding GATA3) and *Calca* (encoding CGRP). Bottom right: cell type annotations. (**D**) Significance (Bonferroni-corrected *p*-value, dot color) and effect size (dot size) of overlap between the program genes (topics, columns) and Gene Ontology (GO) categories (rows) for the topics in B. (**E–G**) Program QTLs. (**E**) Significance (*y-*axis, –log_10_(*P*-value)) of association of each genomic locus (*x*-axis) with the cell score of a given topic (top left label). Dotted red line: genome-wide significance *p*-value = 0.05 (permutation-derived significance; Bonferroni corrected). Blue: significant associations. (**F**) A proportion QTL overlapping *Sstr4*. Gene models (top), conserved regulatory elements (CRE) and LOD scores (*y*-axis) of QTL effects in a locus including *Sstr4* (red vertical stripe: program QTL) and associated with topics 1 (blue) and 3 (pink). (**G**) Shared and specific genes in program QTLs associated with topics 1 and 3. (**H**) LOD score (*y*-axis) along each of three QTLs (*x*-axis) associated with topic 8 in the DO mouse analysis (bottom, black) and for the founder strain QTL effect coefficients inferred from the DO mapping model (top, colors). Red dashed horizontal line: genome-wide significance *p*-value = 0.05 (permutation-derived significance, Bonferroni corrected). Black dashed vertical line shows alignment between LOD peak in DO mouse analysis and the founder strain QTL effect coefficient.

We next found 30 “program QTL” associated with at least one of 14 ILC topics, with some QTL impacting multiple programs, and some programs associated with multiple QTL (**Fig. 3E, Fig. S3B, Methods**). For example, the program QTL associ-ated with topic 1 (ILC1 and ILC2) and 3 (CCR6^high^ LTi) over-lap homeostasis regulators (**Fig. 3G**). These include a QTL in chromosome 2 associated with both topic 1 and topic 3, with the strongest signal overlapping *Sstr4*, encoding somatostatin receptor 4, which in the intestine regulates the expression of the peptides CGRP, SP, SOM, SSTRs, and neuron innervation or nociception^72^ (**Fig. 3F**). *Sstr4* is not expressed in any ILC sub-sets but is expressed in multiple mouse enteric neuron subsets^73^ (**Fig. S3C**). Note that *Cd93*, is also proximal to (but does not overlap) the same QTL that overlaps *Sstr4*, is also an eGene in a *cis*-QTL in ILC2s, ILC3s^74^ and LTis and may also be the putative causal gene in this locus. In another example, ILC3/ LTi topic 8, which includes *Il22, Rorc*, and *Jun*, is associated with 3 QTL on chromosome 3, 5, and 11 (**Fig. 3E**), each likely to im-pact this program in different founder strains, based on founder strain QTL effect allele contribution coefficients inferred from the mapping model (**Fig. 3H, Methods**).

### Pleiotropic QTL impact multiple ILC traits as well as serum cytokine levels

A substantial number of QTL may be pleiotropic in that they are associated with multiple ILC traits, within a category (eQTL, proportion QTL or program QTL) or across categories. Specifically, 108, 897, and two QTL were associated with more than one expression, proportion, or gene program trait, respectively; and 328, 24, and 12 QTL were associated with both expression and proportion, expression and gene programs, or proportions and gene programs, respectively (**Fig. 4A–C**). (As above, we note that some of the traits are likely not independent.)

**Fig. 4.**
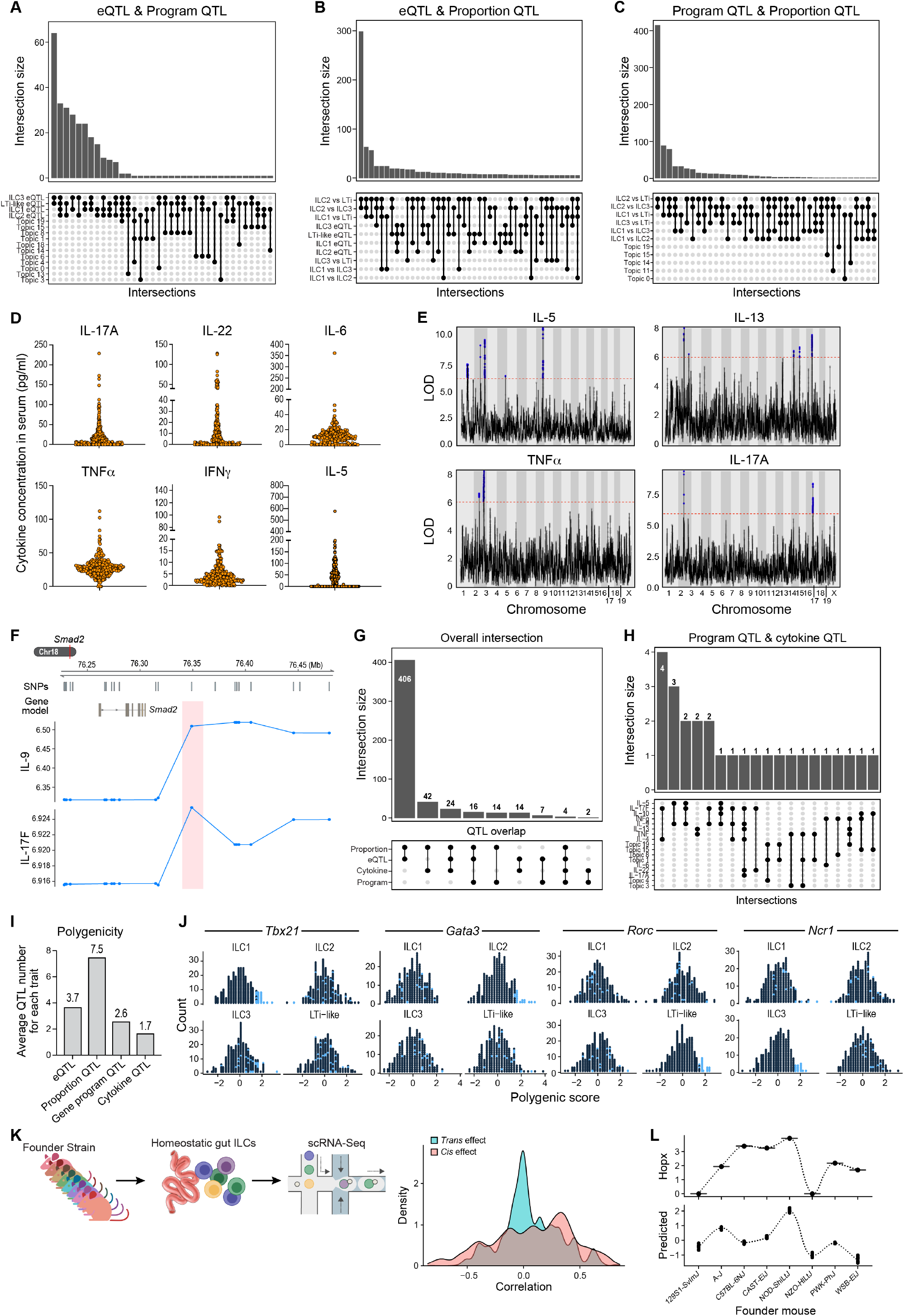
Pleiotropic QTLs impact multiple ILC traits and serum cytokine levels and polygenic architecture of ILC traits enriched for specific path-ways. (**A–C**) Shared expression, proportion and program QTLs. Number of loci (*y*-axis) shared between pairs of trait classes (*x*-axis, as marked on bottom): expression and program (**A**), expression and proportion (**B**), and program and proportion (**C**). (**D**) Variation in peripheral serum cytokine levels across 261 DO mice at homeostasis. Cytokine level (*y*-axis) for each cytokine (panel) in each mouse (dot). (**E**) Cytokine QTLs. LOD score (*y*-axis) for association of each genomic locus (*x*-axis) with a serum cytokine’s level (labeled on top). Dotted red line: *p*-value = 0.05 (Bonferroni corrected). Blue: significant associations. (**F**) A cytokine QTL proximal to *Smad2*. Gene models (top), conserved regulatory elements (CRE) and LOD scores (*y*-axis) of QTL effects in a locus including *Smad2* (red vertical stripe: cytokine QTL) and associated with variation in levels of serum IL-9 and IL-17F. (**G–I**) Cytokine QTLs overlap with ILC trait QTLs. Number of loci (*y*-axis) shared between pairs of trait classes (*x*-axis, as marked on bottom) across expression, program, proportion and cytokine QTLs overall (**G**), or specific cytokine and program QTLs (**H**). (**I**) Polygenicity of expression, proportion, program and blood cytokine traits. Mean number of QTLs (*y*-axis) associated with each trait category (*x*-axis). (**J**) Polygenic, cell type specific association for expression of key ILC TFs. Distribution of polygenic scores (*x*-axis; based on correlation between gene alleles and expression level (**Methods**)) for each of four key TF genes in each ILC subset. Blue: mice with high PRS score for the TF’s expression in its cell type. (**K, L**) Stronger heritability of *cis*-eQTL effects based on analysis of founder strains. (**K**) Distribution of correlation coefficient between predicted and observed gene expression for each founder strain allele for *cis*- (blue) and *trans*-(red) eQTLs. (**L**) Measured (top, from founder mice) and predicted (bottom, from DO mice analysis) expression (*y*-axis) of *Hopx* for each founder strain allele (*y*-axis) in ILC1s.

To explore the broader impact of such pleiotropic QTL we examined their association to peripheral serum cytokine levels. We collected the serum from each of another 261 DO mice that were at homeostasis, and measured the levels of multiple cytokines that are associated with ILC effector and regulatory functions, including IL-2, IL-4, IL-5, IL-6, IL-9, IL-10, IL-13, IL-17A, IL-17F, IL-22, IFN-γ and TNF-α^3,7,75^ (**Methods**). There was substantial variation in cytokine levels across the DO mice, and most cytokines grouped in one of two clusters by the cor-relation in their levels across mice (**Fig. 4D, Fig. S4A, B**).

QTL mapping identified 41 cytokine QTL (**Fig. 4E, Fig. S4C, Methods**), including two strong loci (genome-wide LOD > 6) in chromosome 2 associated with multiple cytokine QTL: one overlapping the HoxD cluster and another overlapping *Sec23b*, a gene critical for cytokine secretion from T cells^76^. In another example, *Smad2*, a regulator of Th17 differentiation and intestinal function^77–79^, is proximal to a QTL associated with the levels of the pro-inflammatory cytokines IL-9 and IL-17F, which are related to intestinal barrier injury^80^, and correlated across the mice (**Fig. 4F, Fig. S4A, B**).

Importantly, 34 of the 41 cytokine QTL overlapped with ILC QTL of either subset-specific expression (15 QTL), program (5 QTL) or cell proportion (14 QTL) (**Fig. 4G, H, Fig. S4D, E**). For example, a QTL overlapping *Bcl2* and *Kdsr* was associated with both serum IL-5 levels and topic 19, active in ILC1 and ILC2s (**Fig. S4F**). Another QTL associated with variation in both *Il18r1* and *Hspa14* expression in intestinal ILC1s and serum levels of the ILC1 signature cytokine IFNγ^81^ (**Fig. S4G**) overlaps *Ptpn5* (STEP), an NKT-expressed gene involved in the differentiation and activation of γ*δ* T cells (which produce IL-17A and IFNγ^79,82^) (**Fig. S4H**), within a gene dense region with multiple itch receptor genes (Mrgprs) that are expressed in sensory neurons and mediate neuron-immune crosstalk in regulating mucosal homeostasis^83^.

### Polygenicity and heritability of ILC traits

Generally, ILC traits — including cell type specific expression, subset proportions and gene programs — were polygenic, associated, respectively, with 3.7, 7.5, and 2.6 QTL per trait on average, whereas peripheral cytokine levels were associated with fewer QTL (1.7 on average) (**Fig. 4I**). Interestingly, while we did not identify individual significant eQTL for the level of key known ILC TFs (e.g., *Tbx21* (ILC1), *Gata3* (ILC2), *Rorc* (ILC3 and LTi))^6^, we did find a polygenic association based on cell type specific “polygenic risk scores” (PRS) for these traits, with continuous, normally distributed, and evident ILC subtype-specific patterns (**Methods**). For example, DO mice bearing polygenic alleles positively affect *Tbx21* expression solely in ILC1s (high “PRS”, **Fig. 4J**). Interestingly, we found a positive polygenic effect on *Rorc* expression in LTis but not in conventional ILC3s, and a polygenic association for *Ncr1* (rather than *Rorc*) in LTi-like ILC3s (**Fig. 4J**). This is consistent with previous studies showing that *Ncr1* distinguishes classic ILC3s from LTis especially in the intestine^84,85^.

We further leveraged the founder strains to estimate the heritability of expression variation from individual QTL, by profiling ~80,000 intestinal ILCs from the eight founder strain mice and comparing the predicted founder strain QTL effect coefficients at variant loci from DO mice to the observed effects in the founder strain mice (**Methods**). Notably, genes whose expression is associated with *cis*-eQTL had overall higher correlation between predicted and observed expression than those with *trans*-eQTL associations, similar to reported observations for pQTL in the DO mice^18^ (**Fig. 4K**). For example, *Hopx* expression levels in ILC1s predicted by *cis*-eQTL regions from the DO mice were significantly correlated *(r* = 0.22; *P* < 10^−5^) with the observed levels across the founder strains (**Fig. 4L**).

### QTL overlapping only non-ILC expressed genes are prevalent *trans*-regulators of ILC genes, programs, and proportions

Both analysis of individual loci and global analysis of QTL suggested that many QTL may impact ILC traits in a non-cell intrinsic manner. For example, a QTL overlapping *Il22ra1* is associated with ILC3 vs. ILC2 proportions, but *Il22ra1* is expressed in intestinal epithelial cells^86^, whereas its ligand, IL-22, is secreted by ILC3s, suggesting a potential circuit that impacts the balance of ILC3 vs. ILC2 (**Fig. 5A**). As noted above, a QTL associated with *Il18r1* and *Hspa14* expression and serum IFNγ level, includes PTPN5, expressed in non-ILCs in the intestine (NKT cells and CD8^+^ T cells; **Fig. S4H**), and multiple itch receptor genes (Mrgprs), expressed in enteric neurons and important for neuro-immune interaction in mucosal tissues^83^. Other non-ILC expressed genes in *trans-*QTL included *Muc2* (overlapping a *trans* eQTL that is associated with *Stub1* and *Ift20* expression in ILC1s, **Fig. 1G**), and *Fam21* (overlapping a proportion QTL associated with proportion of ILC1 vs. ILC2) (**Fig. S2C**).

**Fig. 5.**
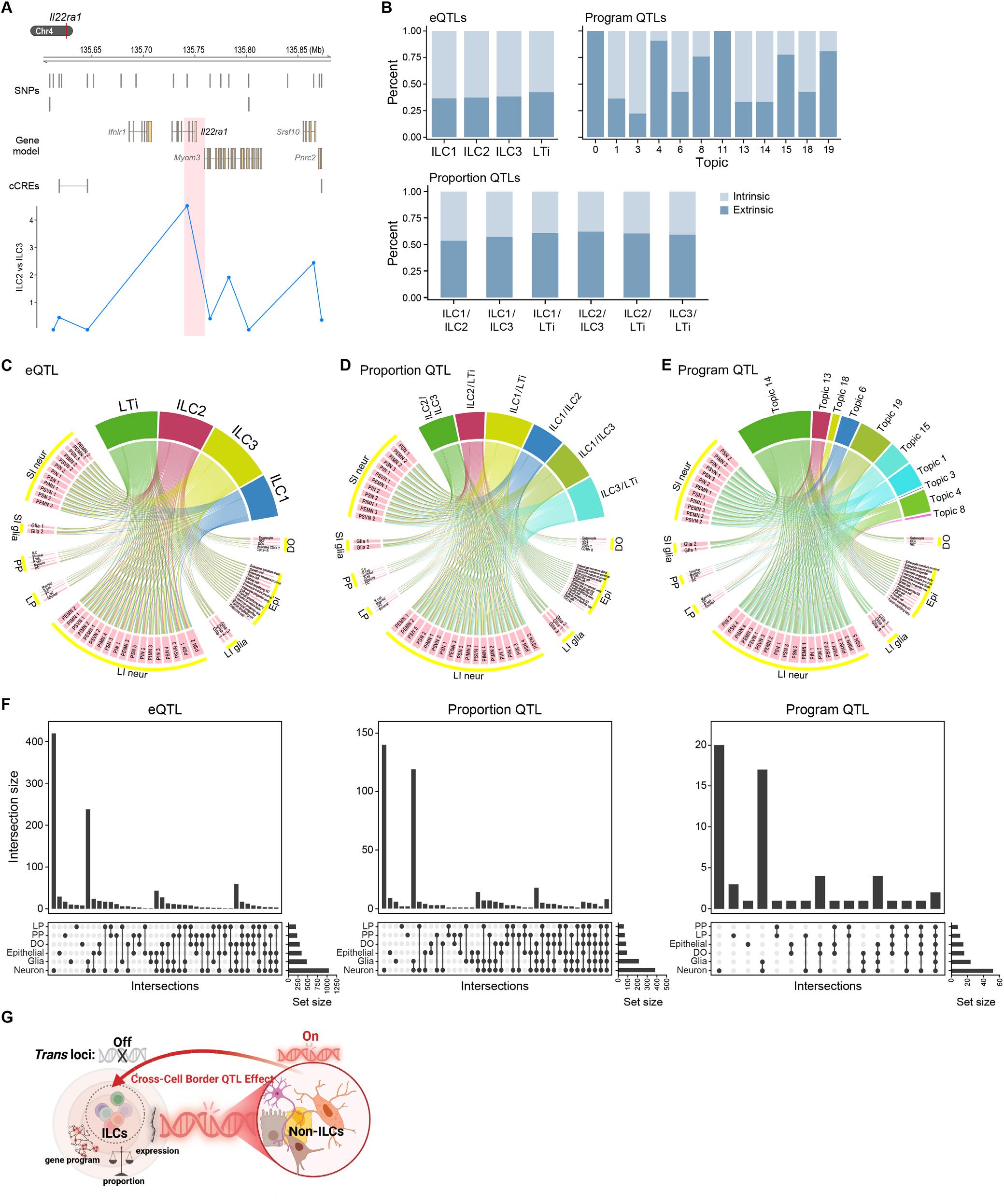
QTLs with non-ILC genes are prevalent *trans*-regulators of ILC genes, programs and proportions. (**A**) QTL overlapping *Il-22ra* associated with ILC3/ILC2 proportion. Gene models (top) and LOD scores (*y*-axis) of QTL effects in a locus including *Il-22ra* (red vertical stripe: proportion QTL) associated with the relative proportion of ILC3/ILC2. (**B**) Many *trans* QTLs do not overlap any ILC expressed genes. Fraction of *trans* QTLs (*y*-axis) overlapping no ILC expressed genes (dark blue) or overlapping at least one ILC-expressed gene (light blue). (**C–F**) QTLs with no ILC expressed genes overlap many enteric neuron, glia and epithelial cell expressed genes. (**C–E**) Genes overlapping expression (**C**), proportion (**D**) or program (**E**) QTLs with no ILC-expressed gene in each trait (top bars) expressed (>25%) in different cell types from the mouse intestine in different scRNA-seq datasets. LP: lamina propria10; PP: Peyer’s patches^10^, SI neur: enteric neuron in small intestine^73^, SI glia: enteric glia in small intestine^73^, LI neur: enteric neuron in large intestine^73^, LI glia: enteric glia in large intestine^73^; Epi: Intestinal epithelial cells^20^; DO: non-ILC scRNA-seq in this study. (**F**) Number of genes (*y*-axis) from QTLs overlapping no ILC-expressed genes expressed (>25%) in each combination of cell types (*x*-axis and bottom rows) from mouse intestine scRNA-seq studies.

Indeed, to the best of our analysis, on average, 48% of *trans* QTL overlapped no gene expressed in ILCs (37%, 59%, 61% for expression (*trans*), proportion, and program QTL, respectively) (**Fig. 5B**). We analyzed the genes overlapping these loci (within +/– 50 kb of the most significant SNP) for expression in other cell types based on scRNA-seq from the mouse small intestine, including our DO mice (non-ILCs captured in our gates; **Fig. 1B**), immune cells from the lamina propria and Peyer’s patches at homeostasis^10^, the small intestine epithelium^20^; and the enteric nervous system^73^ (all generated in our lab). The genes overlapping these QTL were expressed in a broad range of non-ILC cells (**Fig. 5C–E, G**), most prominently in enteric neurons and intestinal glia (**Fig. 5F, G**, *P* < 10^−16^, Fisher’s exact test). For example, 89.3% of genes overlapping *trans* eQTL with no ILC-expressed genes are expressed in neuronal cells (36% uniquely in neurons), 41.3% in glia, and 27% in epithelial cells (**Fig. 5F, G**).

### QTL identity cross cell type epithelial-, phagocyte- and neuron-ILC interactions and feedback in intestinal tissue

Analysis of specific QTL overlapping only non-ILC genes suggests causal cell–cell interactions in tissues^4,6^. We highlight a few illustrative examples, spanning epithelial-, phagocyte- and neuron-ILC tissue circuits from our analysis.

In some cases, a QTL for an ILC trait overlaps or is proximal to a cytokine gene expressed in an immune or epithelial cell type, while its receptor is expressed on an ILC subset. For example, a *trans* eQTL impacting the ILC2-expression of *Dap3, Myl12b*, and *Tmsb4x* overlapped *Ccl17* (**Fig. 6A**), a chemokine known to be secreted by dendritic cells (DCs), macrophages and monocytes to mediate tissue inflammation acting through a DC-CCL17-Th2-IL-4 axis^87–89^. *Ccl17* is not significantly expressed in ILCs, but its receptor *Ccr4* is selectively expressed in ILC2s (**Fig. 6B, Fig. S5G**). This suggests the possibility of a circuit from DC-CCL17 to ILC2s-CCR4 impacting on ILC2 gene expression.

**Fig. 6.**
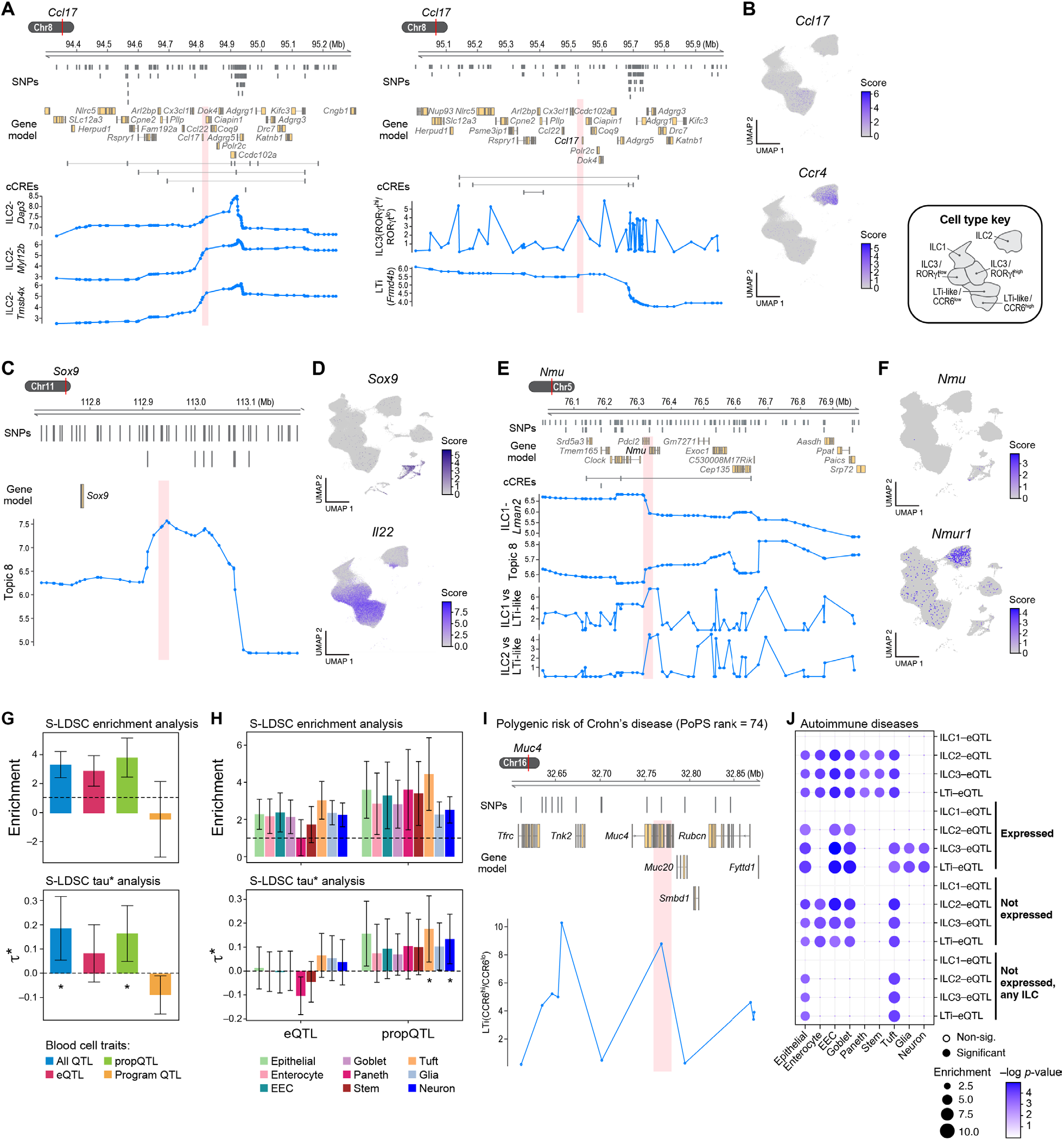
Intercellular *trans*-regulation of ILC traits reveals epithelial-, phagocyte- and neuron-ILC circuits. (**A, B**) DC-expressed *Ccl17* associated with ILC2 expression. (**A**) Gene models (top) and LOD scores (*y*-axis) of QTL effects in a locus overlapping *Ccl17* and other genes (red vertical stripe: QTL) and associated with ILC2-expression of *Dap3, Myl12b*, and *Tmsb4x*. (**B**) UMAP embedding of ILC profiles (as in Fig. 1C) colored by expression of *Ccl17* (top) and its receptor *Ccr4* (bottom). Right: Cell type key. (**C, D**) Epithelial-ILC feedback loop inferred from *Sox9*–*IL22*-expressing topic 8 association. (**C**) Gene model (top) and LOD scores (*y*-axis) of QTL effects in a locus overlapping *Sox 9* and associated (red vertical stripe: program QTL) with topic 8 (high-scoring genes: *Rorc, Jun, Fosl2, Il22, Cd69, Il7r*). (**D**) UMAP embedding of ILC profiles (as in Fig. 1C) colored by expression of *Sox9* (top) and *Il22* (bottom). (**E, F**) ENS-expressed *Nmu* associated with multiple ILC traits. Gene model (top) and LOD scores (*y*-axis) of QTL effects in a locus overlapping *Nmu* and additional genes and associated (red vertical stripe: QTL) with multiple ILC traits (bottom). (**D**) UMAP embedding of ILC profiles (as in Fig. 1C) colored by expression of *Nmu* (top) and its receptor *Nmur1* (bottom). (**G**) Human orthologs of proportion QTLs are enriched for autoimmune disease heritability. Sc-linker meta-analyzed heritability enrichments (top panel) and standardized effect sizes (bottom panel) across 11 relatively-independent autoimmune diseases for QTL overlapping genes from all QTLs (blue), eQTLs (red), proportion QTLs (green) and program QTLs (yellow), using enhancer-gene links aggregated across all tissues (**Methods**). All results are conditional on the 87 baseline-LD (v2.1) annotations. (**H–J**) Human orthologs of proportion QTLs not expressed in any ILCs but expressed in tuft cells or neurons are enriched for autoimmune disease heritability. (**H**) sc-linker meta-analyzed heritability enrichments (top panel) and standardized effect sizes (bottom panel) across 12 relatively independent autoimmune diseases for expression and proportion QTL overlapping genes that are not expressed in any ILCs but expressed in one or more of nine other cell types (color code). Gene models (top) and LOD scores *(y*-axis) of QTL effects in a locus overlapping *Muc4* (red vertical stripe: proportion QTL) and other genes and associated with the relative proportion of CCR6^high^LTi/CCR6^low^ LTi. (**J**) Significance (–log_10_ (*P*-value), dot color) and magnitude (dot size, E-score) of sc-linker meta-analyzed heritability enrichments across 12 autoimmune diseases for eQTL overlapping genes with 16 different characterizations of expression across 4 ILC cell sub-types (rows, Methods) and expressed in 9 different non-ILC cell types (columns).

In other cases, a gene with non-ILC, cell type-specific expression resides in a QTL that causally affect ILC traits, and the ILC traits in turn impact the source cell of the gene, potentially closing a tissue feedback loop. For example, a QTL associated with ILC3/LTi topic 8 is proximal to only one gene, *Sox9*, a TF in intestinal epithelial cells, which regulates Paneth cell regeneration and intestinal homeostasis^90,91^, and is only expressed in our data in enterocytes (**Fig. 6C, D, Fig. S5G, H**). The topic 8 program includes *Rorc, Il-22, Rora*, and *Il7r*, which are key regulators for Th17-like responses, and in turn affect enterocyte function^86,92^, thus closing a putative feedback loop in the tissue modulating intestinal homeostasis: from *Sox9* (in enterocytes) to (ILC3/LTi) topic 8 (including *Il-22*), to IL22 secretion back to enterocytes^78,86,93^.

The cross-cell QTL effects also re-discover and complement known findings on neuron-ILC interactions. Neuronally expressed neuropeptides, such as neuromedin U (NMU), substance P (SP) and vasoactive intestinal peptide (VIP) are all encoded by genes overlapping QTL associated with various ILC traits^13^. Specifically, the *NMU* gene overlaps a pleiotropic QTL associated with ILC1/LTi and ILC2/LTi proportions, ILC1 *Lman2* expression, and topic 8 (**Fig. 6E**). We and others have previously shown a neuron-NMU-ILC2 axis in the small intestine^42,94^, where *Nmu*, highly expressed by cholinergic enteric neurons, regulates the ILC2 response to worm infection^71^. *Nmu* is specifically expressed in enteric neuro subsets (e.g., PSN1, PEMN273) and not in ILCs, and its receptor *Nmur1* is highly expressed in ILC2^94^ (**Fig. 6F, Fig. S5G, H**). In another example, a QTL associated with CCR6^high^/CCR6^low^ LTi proportions and *Sec61b* and *Rpn2* expression in LTis, is proximal to *Tac1*, the gene encoding Substance P (SP), a neuropeptide and neuromodulator (**Fig. S5A, B, G and H**). SP is known to act on mast cells to promote the release of inflammatory mediators in the skin^95^ and to regulate smooth muscle contractility, epithelial ion transport, vascular permeability, and immune function in the gastrointestinal tract in IBD pathophysiology^96–98^. This suggests an SP-ILC axis that can impact *Rpn2* and *Sec61b* and LTi cell state at intestinal homeostasis. Finally, both the neuropeptide VIP (expressed in neurons) and one of its receptors *Vipr2* (VPAC2, expressed on LTis and ILC2s) overlap different QTL, associated with proportion of RORγt^high^ ILC3/RORγt^low^ ILC3s (both), peripheral IL-2 level (*Vip*) and LTi expression of *Ablim1, Rpl35*, and *Slx1b* (*Vipr2*) (**Fig. S5C–H**). This is consistent with studies showing that VIP, binding to VPAC2 (*Vipr2*) on ILC3s (ILC3 and LTi), stimulates the production of IL-22, critical in the maintenance of immune homeostasis and metabolic processes at intestinal barrier^99,100^.

### Gene ortholog sets defined by DO expression and proportion QTL are enriched for human autoimmune disease risk heritability

Finally, we explored the relevance of our findings to human disease, by assessing the contribution of human orthologs of genes overlapping different QTL categories — expression, proportion or program QTL — to heritability of relevant human traits, such as autoimmune disease risk. To this end, we identified human orthologs for the genes overlapping different sets of QTL (defined as genes within +/– 50 kb of the most significant SNP) in the DO mice and integrated them with genome-wide association studies (GWAS) for common variants associated with 12 immune-related diseases and 7 blood cell traits (genetic correlation: *r*_*g*_ < 0.9)^101^. We linked genes to human genetic variants using Roadmap^102^ and Activity-By-Contact^103,104^ enhancers and assessed disease heritability using sc-linker^25^, conditioning all heritability results on 86 baseline-LD (v2.1) annotations (**Methods**). We assessed all genes in a QTL category, as well as subsets that included only ILC-expressed or non-ILC expressed genes.

Overall, genes overlapping expression and proportion QTL across ILC subsets were enriched in heritability in both auto-immune diseases (**Fig. 6G**) and blood cell traits (**Fig. S6A**). The lack of heritability enrichment in program QTL genes may be due to limited power given the smaller number of QTL (and overlapping genes). Trait-level heritability varied across diseases and QTL categories (**Fig. S6A**). Genes overlapping proportion QTL also showed conditionally informative disease heritability signals for autoimmune diseases, contingent on the baseline-LD annotations (τ* = 0.17, *p*-value = 0.001; **Fig. 6H**, bottom, asterisk), but not in the meta-analysis across blood cell traits (**Fig. S6H**, bottom). Moreover, genes overlapping proportion QTL that were not expressed in any ILCs but expressed in at least one of nine other gut cell types also showed conditional disease heritability signal (**Fig. 6H, Fig. S6C**). Among those, genes overlapping proportion QTL that are expressed in tuft epithelial cells (but not in ILCs) had the highest autoimmune heritability enrichment (4.4x, *p*-value = 3*10^−6^), as well as significant conditional disease information τ* = 0.175, *p*-value = 0.006), as did proportion QTL genes expressed in neurons (but not in ILCs) (τ* = 0.133, *p*-value = 0.003) (**Fig. 6H**). As a secondary result, proportion QTL overlapping genes expressed in tuft, glia and neurons, irrespective of their expression in ILCs, showed conditional autoimmune disease signal (**Fig. S6C**). One example is *MUC4*, a gene overlapping a LTi (CCR6^high^/CCR6^low^) proportion QTL, which is not expressed in any ILC subset but expressed in mouse neuron and glia and is a top 100 gene for the polygenic risk of Crohns’ disease^105^ (PoPS rank^105^ = 74) (**Fig. 6I**). *Muc4* overexpression had previously been implicated as a driver of Crohn’s disease^106^ and shows eQTL colocalization with mean corpuscular volume in GTEx brain cerebellum^107^.

Genes overlapping specific expression QTL for different ILC subsets, either expressed in ILCs or only expressed in other gut subsets also had enriched heritability and standardized effect sizes (τ*) in a meta-analysis across autoimmune diseases (**Fig. 6J, Fig. S6D**). For example, ILC3 and LTi eQTL-overlapping genes expressed in glia and neurons (as well as in corresponding ILC subsets) showed heritability enrichment signal for autoimmune diseases (**Fig. 6J**). In addition, genes overlapping ILC subset eQTL with expression in other gut cell subsets that themselves are not expressed in the same ILC subset (but can be expressed in other ILC subsets) had 1.7x stronger enrichment in autoimmune disease than those not expressed in any ILCs (**Fig. 6J**). Finally, genes overlapping ILC1 eQTL had 2.2x higher disease enrichment signal compared to the other ILC subset QTL in a meta-analysis across blood cell traits; in contrast, these genes did not show significant autoimmune disease signal (**Fig. 6J, Fig. S6E, F**).

Overall, this analysis suggests that genes from expression and proportion QTL from DO mice point to processes with relevance to heritability of human immune diseases as well, both for genes expressed in ILCs and for ones specifically expressed in other cell types, especially neurons and tuft epithelial cells.

## DISCUSSION

With unprecedented advances in the discovery of genetic variation and the assembly of the human cell atlas, the next challenge is to characterize the associations between genetic variation and phenotype at scale, and their underlying mechanisms^108^. Here, we combined the DO mice model system with scRNA-seq to identify QTL associated with intestinal ILC traits, including subtype-specific eQTL, program QTL and cell subset proportion QTL.

Most notably, while most QTL studies focus on cell intrinsic effects, our analysis highlights the power of QTL combined with single cell genomics to also recover likely cell extrinsic effects, where the genetic variant in a locus overlapping a gene expressed in one cell type in a tissue is associated with a phenotype in another cell type in that tissue. In many loci, the strongest signal (or the entire locus) spans genes that are not expressed in ILCs, but are expressed in other cells in the tissue, such as enteric neurons, intestinal epithelial cells, and intestinal glia. These allowed us to decipher putative cross cell type causal interactions in tissue.

A very recent study^27^ also highlighted genetic evidence for cross cell type genetic effects between immune cells. Notably, however, those were PBMCs and thus not collected in a tissue context, and the traits were only cell proportions (based on marker protein expression) which limit any potential mechanistic implications. Nonetheless, they suggest that this approach could generalize across systems.

Some of the functional circuits we recovered are well validated, including the crosstalk between ILCs and neurons through neuronal peptides NMU, CGRP, SP, or VIP, but our analysis sheds light on the specific impact on ILCs. In other cases, the circuits we uncovered have not been previously characterized. Some traits are associated with QTL overlapping genes expressed in multiple other cell types. For example, topic 8, expressed in ILC3s and LTis and including key signature genes encoding cytokines and TFs, such as *Il22, Rora* and *Rorc*, is associated with QTL overlapping genes expressed in phagocytes, enterocytes, and neurons, which may affect ILCs directly through matching receptor/ligand interactions or indirectly through additional cells and global functions in the tissue.

Our heritability analysis shows that DO-defined QTL may be relevant for human autoimmune disease, with enriched heritability observed for both expression and proportion QTL genes overall, for specific subsets, and for proportion QTL genes expressed only in tuft cells or neurons. More generally, our approach could be further useful in human genetics of disease, where uncovering causal cell–cell circuits in tissue is far more challenging than in animal models. To date, most single cell profiling studies of human genetic variation have focused on cells (e.g., PBMCs or iPSCs) and only few on complex tissue. As single cell and spatial genomics can now be applied at scale to larger cohorts^109,110^, it is likely that similar cross-cell type genetic effects can be recovered as well, but they may require studying associations in *trans* to traits like cell proportions or gene programs. Moreover, extending studies like ours to profile entire tissue ecosystems (rather than a single cell type here), should allow recovering more complex circuitry and possibly mediating cell types for indirect interactions, to help decipher the mechanistic basis of disease.

## Limitations of the study

Linkage mapping in DO mice has lower resolution than human mapping studies. Thus, it is sometimes challenging to pinpoint the likely gene from a multi-gene locus (especially if genes are expressed in both ILCs and non-ILCs), and each QTL region must be interpreted with care. Our study focused on just one cell type (ILCs): The interactions between cells across the tissue could be substantiated further by both full tissue analysis of the entire cellular ecosystem, and ideally in a spatial context. These would allow better determination of direct vs. indirect cell–cell effects. Finally, our study was limited to mice (and ortholog-based projection to human). Future analysis of human tissues will help determine the relevance of this approach in human genetics.

## METHODS

### Animals

Diversity outbred (DO) mice (Jax stock #009376) and 8 founder strain mice (C57BL/6J, 129S/SvlmJ, PWK/PhJ, NOD/ ShiLtJ, A/J, NZO/HlLtJ, CAST/EiJ, WSB/EiJ) were purchased from The Jackson Laboratory. Mice were housed under specific pathogen-free conditions at the animal facility at the Broad Institute. All mouse work was performed in accordance with the Institutional Animal Care and Use Committees (IACUC) and with relevant guidelines at the Broad Institute and Massachusetts Institute of Technology, under protocol 0055-05-15. 7–9-week-old female mice were used for all experiments.

*Rorc*-cre (Jax 022791) mice were originally obtained from the Jackson Laboratory. *Rbpj*^flox/flox^ mice were generously provided by Professor Huan Han^111^. *Rbpj*^flox/flox^ and *Rorc*-Cre mice were bred to obtain the *Rbpj*^fl/fl^ *Rorc*-Cre strain. Mice were housed in specific pathogen-free conditions and were used and maintained in accordance of Institutional Animal Care and Use Committee guidelines of Westlake University. Both male and female mice were used at 8–10 weeks unless otherwise indicated.

### Isolation of cells from the lamina propria for flow cytometry

Intestines were opened longitudinally and washed in ice-cold PBS. Epithelial cells and intraepithelial lymphocytes were removed by incubating the tissues for 20 min at 37°C with constant stirring at 350 rpm in extraction buffer (5% fetal bovine serum, 1 mM DTT (ThermoFisher Scientific), 5 mM EDTA (ThermoFisher Scientific) in RPMI 1640 medium (ThermoFisher Scientific)). Tissues were minced and incubated in digestion buffer (5% fetal bovine serum, 50 μg/ml DNase I (Sigma Aldrich), 1 mg/ml Collagenase Type VIII (Sigma Aldrich), 2 mM CaCl2 (Sigma Aldrich), 5 mM MgCl2 (Sigma Aldrich) in RPMI 1640 medium (ThermoFisher Scientific)) for 30 min at 37°C with constant stirring at 550 rpm and smashed through a 40 μm sterile strainer. Mononuclear cells were enriched on a discontinuous (37% and 70%) Percoll gradient (Cytiva). Red blood cells were removed using ACK lysis buffer (Lonza) at room temperature (RT) for 2 min.

### Flow cytometry

Cells were washed and prepared as single-cell suspensions in MACS buffer (pH 7.4; PBS plus 1% FBS and 5 mM EDTA), followed by blockade of non-specific binding by incubation for 10 min on ice with 25 μg/ml 2.4G2 (BD) before the addition of specific antibodies to cell-surface antigens. To detect intra-cellular proteins, cells were further fixed and permeabilized for 30 min at 25°C with a Foxp3 Staining Buffer Set according to the manufacturer’s protocols (eBioscience). Non-specific binding in fixed cells was blocked by incubation for 10 min at 25°C with 25 μg/ml 2.4G2 (BD). Cells were typically stained at 4°C for 30 min. Antibodies and staining reagents were from eBioscience unless otherwise stated: Alexa Fluor 700 anti-CD45, Allophycocyanin (APC) anti-CD3e, APC anti-Ly6G/Ly6C, APC an-ti-CD19, APC anti-CD11c, APC anti-CD11b, APC anti-TCRγ/*δ*, APC anti-TCRβ, APC anti-CD5, APC anti-FcεRIα, APC an-ti-NK1.1, BV605 anti-CD90.2 and PE-cy7 anti-NKP46, Fixable Viability Dye eFluor™ 780 (ThermoFisher), phycoerythrin (PE) anti-RORγt (ThermoFisher), FITC anti-KLRG1 (ThermoFisher) and BV421 anti-CCR6 (BD). For intracellular cytokine staining, lamina propria cells were stimulated with Cell Stimulation and Protein Transport Inhibitor Cocktail (eBioscience) in complete RPMI medium (10% FBS, 10 mM HEPES, 1 mM sodium pyruvate, 80 µM 2-mercaptoethanol, 2 mM glutamine, 100 U/mL penicillin, 100 mg/mL streptomycin) for 4 hours at 37°C with 5% CO2. Following stimulation, cells were stained with antibodies for surface markers as mentioned above, and subsequently fixed and permeabilized using Fixation/Permeabilization Solution (ThermoFisher) before staining with AF488 anti-IL17A and PE anti-IL-22 antibodies. Data were collected on Cytoflex (Beckman) and were analyzed with FlowJo software (TreeStar). ILC3s were defined as alive CD45^+^CD90.2^+^RORγt^+^KLRG1^−^ Lineage^−^ (lineage markers: CD3e, CD19, Ly6G/Ly6C, CD11c, CD11b, TCRβ, TCRγ*δ*, CD5, FcεRIα, NK1.1) cells.

### Genotyping with GigaMUGA for DO mice

Tail biopsy samples were obtained from each DO mouse and sent to Neogen (https://genomics.neogen.com/en/mouse-universal-genotyping-array) for isolation of genomic DNA, genotyping and SNP identification using the Mouse Universal Genotyping Array (MUGA) microarrary. The MUGA series contains a 143,259-probe Illumina Infinium II array for the house mouse (*Mus musculus*). Most of GigaMUGA is optimized for genetic mapping in the DO mice population, which is highly informative across a wide range of genetically diverse samples with detailed probe-level annotation and recommendations for quality control^33^. In addition to 141,090 SNP probes, GigaMUGA contains 2,006 copy number probes concentrated in structurally polymorphic regions of the mouse genome.

### Small intestinal ILC isolation and scRNA-seq

Intestines were harvested from 274 DO mice and 16 founder strain mice (2 mice for each strain) and placed in a Harvest Buffer (RPMI1640 with 5% FBS) on ice. Fat and feces were cleaned from the tissue and Peyer’s Patches were removed. The jejunum from each set of 5 DO mice were pooled together, cut into small pieces, and transferred to a 20 ml pre-digestion buffer (PBS+ 5% FBS + EDTA (5 mM) + DTT (1 mM)), 37°C for 30 min. Intestine pieces were transferred to a 25 ml wash buffer (PBS with 5% FBS) and vortexed for 30 s. Tissue was placed into a petri dish, chopped with a razor blade until fairly smooth, and digested in 15 ml digestion buffer (RPMI 1640 with HEPES+ 10% FBS + Cacl2 (1 mM, stock: 0.5 M) + Mgcl2 (1 mM, stock: 0.5M) + 100 µg/mL Liberase (Sigma, cat # 5401119001) + 100 ug/mL of DNAse I) for 25 min at 37°C. 1mM EDTA was added to block Liberase activity. Cells were passed through a 100 µm strainer and spun down at 500 *g* for 5 min. at 4°C. Cells were washed with a wash buffer and pelleted again with 500 *g*, for 5 min at 4°C, then resuspended in 4 ml 40% Percoll, and transferred to an FBS coated 15 ml tube. Cells were spun at 600 *g* at room temperature for 20 min, washed again with wash buffer, and pelleted at 2000 rpm for 5 min at 4°C. Cells were resuspended with FACS buffer and ILCs were sorted through gating on live Lin^−^CD45^+^CD90.2^+^IL-7R^+^. A total of 20,000 cells were loaded into each channel of the 10x Genomics Single-Cell Chromium Controller. Multiplexing five mice into one digestion reduced batch effect made the experiment technically feasible and allowed for loading of more cells in each 10x channel for scRNA-seq.

### Bead-based cytokine detection

Blood was collected from the tail vein of each DO mouse and incubated in BD Microtainer® Blood Collection Tubes (BD) for 30 min at 25°C. Tubes were then centrifuged at 25°C, 1000×*g* for 10 min. The serum fraction, located at the top of the tubes, was carefully collected and stored at –80°C for further analysis. Cytokine concentrations in the serum and ILC culture supernatant were determined using LEGENDplex™ MU Th Cytokine Panel (BioLegend, catalog # 741044) according to the manufacturer’s instructions. Samples were acquired using a Beckman CytoFLEX LX Flow Cytometer and analyzed with the LEGENDplex Software v8.0 (BioLegend).

### Characterization of *Rbpj*’s role in ILC regulation

Total small intestinal CD45^+^ cells from *Rorc*^Cre^*Rbpj*^+/+^ and *Rorc*^Cre^*Rbpj*^fl/fl^ were isolated as detailed in the Small Intestinal ILC Isolation section above. Subsequently, flow cytometry analysis was conducted as described above. Composition and cytokine production, specifically intracellular staining of IL-17A and IL-22, were assessed across ILC subsets. Subsets were identified based on defined markers, including: total ILCs (CD45^+^Lin^−^CD90.2^+^), total ILC3s (comprising conventional ILC3s and LTis, both expressing RORγt), conventional ILC3s (RORγt^+^NKP46^+^CCR6^−^ ILCs and RORγt^+^NKP46^−^CCR6^−^ ILCs), and ILC3-like LTis (RORγt^+^NKP46^−^CCR6^+^ ILCs).

### Quantification and statistical analysis

Statistical analysis was performed with GraphPad Prism software version 8.0. Data are shown as mean ± SEM. Statistical significance was determined by Student’s *t* test or two-way analysis of variance (ANOVA) as indicated (two-tailed, \**P* < 0.05, \*\**P* < 0.01, \*\*\**P* < 0.001, and \*\*\*\**P* < 0.0001).

### Haplotype reconstruction

Haplotype reconstruction values were performed as previously described using the GRCm38 mouse genome reference^112^. GigaMUGA intensity values were used after it was observed the genotype calls can be further sub-clustered to the eight founder strain mice as well as the 28 possible F1 genotype combinations between the founder strains. Samples were then estimated to obtain the highest likelihood of a match to a certain founder genotype. Probabilities were combined with a transition probability between the adjacent markers in the array using a hidden Markov model^113^ that also addresses the multiparent population like the founder haplotype structure. Transition probability parameters were selected so that evidence from approximately four sequential markers is necessary to change the haplotype state corresponding to the founder allele. Moreover, the transition penalty varies depending on whether the current state and next state share a common founder. A dynamic programming algorithm was then utilized to calculate the maximum-likelihood founder assignment for each chromosome.

### scRNA-seq demultiplexing

To assign the DO mouse identity for each cell in a channel (pooled from five mice), the Demuxlet^32^ software was used to relate the DO genotype information (above) and scRNA-seq data to assign a probability of the most likely mouse identification to every cell within the dataset. The highest probability assignment from the model was used to recover the mouse identity. Hyper parameter tuning was implemented to balance quality identification with a high number of assigned cells. The mixing parameter, alpha, was used in a grid search to determine an optimum relative proportion each mouse contributed to the cells in the channel, selecting a uniform probability of 20% representation per mouse sample. Other quality considerations were the base quality of the sequence from scRNA-seq reads (minimum Q20) and mapping quality of scRNA-seq reads to the reference genome (*P* < 0.01, i.e., MAPQ > 20).

### scRNA-seq quality control and processing

CellRanger 3.0.2 (10× Genomics) was used to process sequencing reads. BCL files were converted to fastq format using Cell-Ranger mkfastq. Fastq files were fed into CellRanger count using the mm10 reference genome using default settings. Once each channel was processed, the CellRanger agg command was used to aggregate outputs into one dataset, which was loaded into scanpy v1.11.01 for standard preprocessing and filtering. Cell profiles containing fewer than 500 genes or more than 10 percent mitochondrial reads were removed. Scrublet114 was used in combination with the doublet output from demuxlet to remove droplets with a doublet score >0.20. Approximately 371,654 profiles from 274 DO mice and 16,000 profiles from 16 founder strain mice (2 mice for each strain) were retained for downstream analysis.

### Normalization

Normalization was conducted across all cells after filtering. Total UMI counts per gene were normalized to the counts per 100,000 and log-transformed. Normalized counts were used for all downstream analyses.

### Identification of highly variable genes

A loess regression was fit for each genes, expression variance was re-calculated for each gene after standardization to this fitted line and the top 2,000 highest variable genes were selected for further clustering analysis and visualization.

### Batch correction

A light batch correction was performed as previously described^115^ using the single cell channel as a technical batch with Pegasus, which calculates both mean and variance to adjust each channel to the same scale^116^. After adjustment, channels were aggregated to one dataset for analysis.

### Cell type assignment

Dimensionality reduction was performed with principal component analysis (PCA) on the highly variable genes dataset. The top 50 principal components were used to construct a *k*-nearest neighbor graph (*k* = 20) followed by unsupervised clustering with the Louvain community detection algorithm^116^, at multiple resolutions to obtain the number of clusters. Differential expression testing was performed to find marker genes specific to a given cluster was conducted using a Wilcoxon Rank-Sum test with Bonferroni adjusted *p*-value < 0.001, followed by *post-hoc* expert annotation. Annotations were independently confirmed using SingleR(1.4.1; https://rdrr.io/bioc/SingleR/).

### eQTL analysis

Expression quantitative trait loci (eQTL) analysis was performed using the R/qtl2 package^117^, developed for multiparent populations derived from more than two founder strains, such as the Collaborative Cross and Diversity Outbred mice. The coefficient of variation (CV) (STDV/mean) of a given gene’s expression across all the cells of a given subset in one mouse was used as the quantitative trait in a linear mixed model regression^118^ including a kinship matrix to account for population structure. The kinship matrix was constructed following the “leave-one-chromosome-out” method as previously described^119^. Significance was assessed with a permutation test for analysis using a linear mixed model, as previously described^120^, where rows of the haplotype reconstructions were permuted to keep similar population structure while shuffling the mouse ID.

### Overlap with regulatory regions

All eQTL (*cis* and *trans*) were analyzed for overlap with candidate *cis* regulatory elements (cCRE) in ENCODE (https://screen.encodeproject.org/), based on a +/– 50 kb window around the QTL.

### Proportion QTL

To identify genetic loci associated with variation in the proportion of cell subsets, ILC subset proportions (to all ILCs or to another subset) were used as the quantitative trait and a weighted logistic regression was performed. Each proportion trait was analyzed for efficiency in an aggregated form with weights, with the weight variables encoding the number of cells from each mouse in the data. The aggregated binary outcome used was the sum of an indicator of the cell’s assignment to one or the other category in the proportion. Each variant was used as a predictor for all variants in the array. The top 10 principal components of a PCA of mouse genotypes were included as covariates to account for the variation explained by mice relatedness. Multiple hypothesis testing was corrected by the Bonferroni method.

### Topic modeling

Latent Dirichlet allocation (LDA) was used to identify gene programs (topics) across all ILCs with the genism Python package^121^ applied to the expression matrix. A grid search of topics was performed selecting 20 topics based on the calculated model coherence^122^ of the topics^123^. All genes assigned with a non-zero probability for the topic by LDA were considered members of the topic.

### Program (topic) QTL

The topic scores for each cell were used as a quantitative trait for genetic mapping with R/qtl2 analogous to gene expression with a linear mixed model and the same kinship matrix construction and permutation testing as for eQTL analysis above.

### Cytokine QTL

Cytokine levels were used as a quantitative trait for genetic mapping with R/qtl2 analogous to gene expression with a linear mixed model and the same kinship matrix construction and permutation testing as for eQTL analysis above.

### Polygenic inheritance measurement

Polygenic predictions for major lineage-specific transcription factors and marker genes were calculated at a cell subset level. Scores were constructed following the C+T method by each cell type^124^. Using a given eGene as the trait with the markers (SNPs) as predictors and the kinship matrix to account for relatedness among mice as covariates, the effect sizes for each marker were then summed and normalized to represent the final cell type polygenic score for the eGene of interest. Different thresholds for markers to be included were investigated and a best fit was used to construct the final score.

### Gene module expression analysis

Module scores of genes overlapping QTL (+/–50 kb of most significant SNP) were calculated using the AddModuleScore function in Seurat^125^ for the cell profiles in each of three datasets: (1) the small intestine epithelium^20;69^, (2) immune cells from the lamina propria and Peyer’s patches at homeostasis^10^; and (3) the enteric nervous system^73^. Functional enrichment analyses of the genes were performed using the ClusterProfiler R package^74^.

### Sc-linker analysis

Sc-linker was used for computing a disease heritability enrichment score for a set of genes representing a gene program, defined here by human orthologs of genes that overlap (+/– 50 kb of most significant SNP) different types of QTL (expression, proportion and topic QTL) and have different expression patterns in 4 ILC subsets and in non-ILC cell types. Human ortholog genes were determined by using the getLDS() function in biomaRt^126^. SNPs were linked to genes in the gene program using Roadmap^102^ and Activity-By-Contact^103,104^ enhancers linked to human orthologs. Enrichment scores were computed for SNPs based on heritability enrichment of SNPs obtained from stratified LD score regression (S-LDSC^101,127^). Specifically, for each gene set *G*, a set of probabilistic weights between 0 and 1 was constructed for each SNP based on the confidence of them influencing any gene in *G*. Sc-linker computes heritability enrichment estimates EG =% (h^2^ (G))/%SNP(G) for a gene set *G*, and EALL = %(h^2^ (ALL))/%SNP(ALL) for a gene program representing all genes. Here %(h^2^ (A)) corresponds to the fraction of heritability captured by variants linked to genes in program A, and %SNP(A) corresponds to the fraction of variants out of all common and low-frequency 1000 Genomes^128^ variants that are linked to the genes in program A. Finally, an enrichment score for *A* was computed as EG−EALL, where subtracting EALL controls for the baseline level of heritability enrichment for SNPs that influence any gene (since most SNPs do not influence any genes). *P*-values were obtained for the null hypothesis using a block jackknife procedure^101^.

Sc-linker also computes τ*, a metric of unique disease informativeness of each gene program annotation, conditional on a baseline set of 86 coding, conserved and LD-related annotations (baseline-LD v2.1), which is also less affected by the number of variants annotated by an annotation. τ*(*c*) represents the proportionate change in per-SNP heritability associated to a 1 standard deviation increase in the value of the annotation. For annotation *c*, τ* has the following form, τ*(*c*) = τ(*c*) sd_*c*_ / (*h*_*g*_ ^2^ × *M*). τ(*c*) is the contribution of annotation *c* to the per-SNP heritability of a disease conditional on other annotations. *var* (β_*j*_): = ∑_*c*_*a*_*cj*_τ_*c*_ where *a*_*ck*_ denotes the value of annotation *c* in each SNP *j. sd*_*c*_ is the standard error of annotation *c, h*_*g*_ ^2^ is the total SNP heritability, and *M* is the total number of SNPs on which this heritability is computed (equal to 5,961,159 in our analyses).

## Author contributions

H.P.X., K.G., R.J.X., and A.R. conceived and designed the study. G.C., D. G., K. C. and V. P. contributed to the DO experimental design. H.P.X. performed scRNA-seq of DO mice, M.X. performed scRNA-seq of founder strain mice, and the serum cytokine detection of DO mouse. Y.Z. performed the RBPJ validation experiment. K.L. helped with scRNA-seq of DO mice. H.P.X, K.G., and M.X. conceived and designed the core QTL mapping analysis, with guidance from A.R.. K. L. and D. G. helped with analytical method design. M.X., H.P.X. and K.G designed downstream analyses. K.G. performed most of the core analysis, with help from O.A.. M.X., H.X., K.G., and A.R. performed data interpretation. C.H.L., O.A. and M.X. performed gene module expression enrichment analysis and hypergeometric analysis for QTL linked genes. K.D. designed and performed all sc-linker analyses. M.X., K.G., and A.R. wrote the paper with input from all authors. A.R. and R.X. supervised the work.

## Acknowledgments

We thank Leslie Gaffney for help with figure preparation, Heather Kang for help with figure i llustrations a nd t ext editing, and Alok Jaiswal for early help on analyses. This work was supported by the Klarman Cell Observatory at the Broad Institute and National Natural Science Foundation of China (grant 82325023). A.R. was a Howard Hughes Medical Institute Investigator when this study was initiated. Work done at the Broad Institute and MGH was supported by the National Institutes of Health (RC2 DK135492 and P30 DK043351 to R.J.X.), the Helmsley Charitable Trust, and the Klarman Cell Observatory. Work conducted at The Jackson Laboratory was supported by JAX Nathan Shock Center (grant P30 AG038070).

## Inclusion and Ethics statement

Competing interests The authors declare competing financial interests: A.R. is a co-founder and equity holder of Celsius Therapeutics, an equity holder in Immunitas, and was an SAB member of ThermoFish-er Scientific, Syros Pharmaceuticals, Neogene Therapeutics and Asimov until July 31, 2020. From August 1, 2020, A.R. is an employee of Genentech. R.J.X. is a cofounder of Celsius Therapeutics, Jnana Therapeutics, and a member of the scien-tific advisory board of Magnet Bio medicine, Nestle and Moon-lake Immunotherapeutics.

## Data and code availability

Raw scRNA-seq fastq files and processed gene-level data are archived at Gene Expression Omnibus (GEO) under accession number GSE275485. Code is available as an R package from https://github.com/kdgosik/fasi-domice.

---

**Fig. S1.**
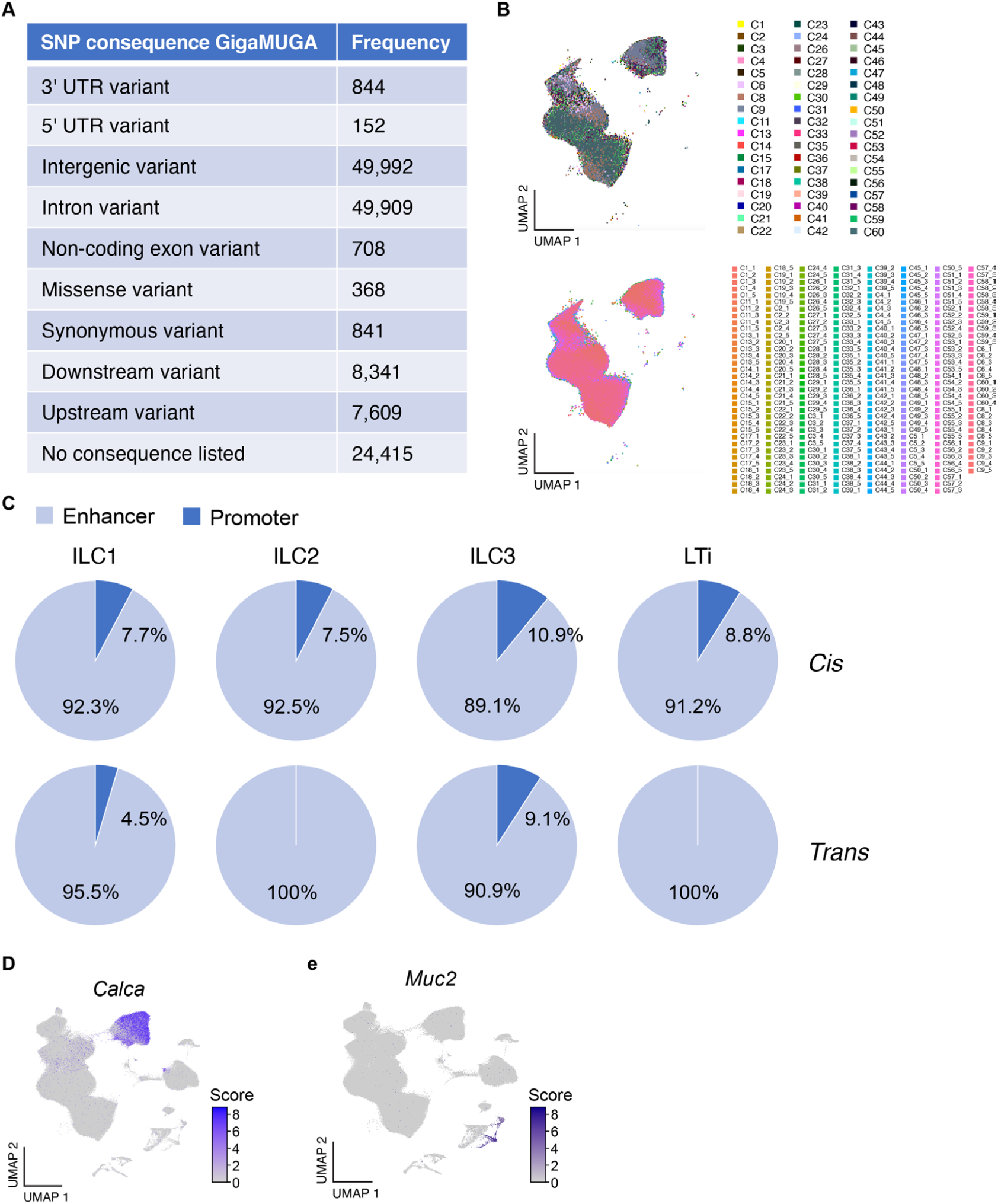
Multiplexed scRNA-seq in DO mice. (**A**) Genomic distribution of genotyped SNPs. Number of SNPs genotyped with GeneSeek Genomic Profiler (GGP) mouse genotyping microarrays (GIGA-MUGA) in each class of genomic elements from the mouse genome. (**B**) Integration of DO mouse scRNA-Seq. UMAP embedding (as in Fig. 1B) of scRNA-seq profiles (dot) colored by batch (top) or mouse (bottom). (**C**) eQTL overlapping regulatory elements are predominantly localized in enhancers. Fraction of *cis* (top) and *trans* (bottom) eQTLs in each cell type present in promoters (dark blue) or enhancers (light blue) out of all eQTLs that overlap (+/– 50 kb) candidate *cis* regulatory elements (cCRE) in ENCODE (https://screen.encodeproject.org/). (**D–E**) Expression of putative causal genes in key eQTLs. UMAP embedding (as in Fig. 1B) of scRNA-seq profiles (dots) colored by expression of *Calca* (**D**) and *Muc2* (**E**).

**Fig. S2.**
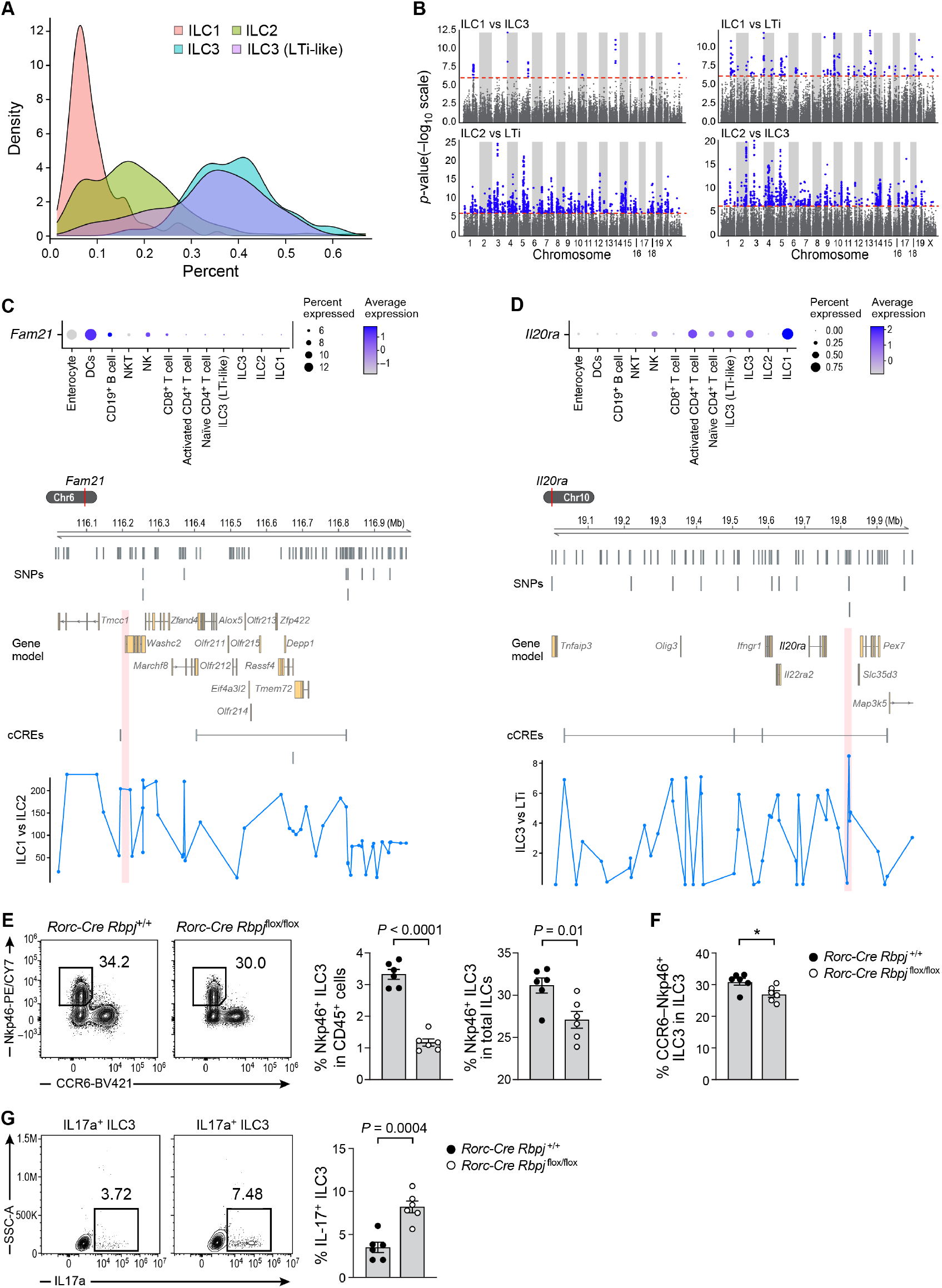
Proportion QTLs. (**A**) Variation in cell type proportions across DO mice. Distribution (density plots) of the fraction of each major ILC subset across mice. (**B**) Proportion QTLs. Significance (*y*-axis, –log_10_(*P*-value)) of association of each genomic locus (*x*-axis) with different cell ratios. Dotted red line: genome-wide significance *p*-value = 0.05 (Bonferroni corrected). Blue: significant associations. (**C, D**) Proportion QTLs in the *Fam21* and *Il20ra* loci. Top: Mean expression (dot color; log(TPM+1)) and proportion of expressing cells (dot size) for *Fam21* (**C**) and *Il20ra* (**D**) across different cell subsets (rows) in DO mice. Bottom: Gene models (top), conserved regulatory elements (CRE) and LOD scores (*y*-axis) of QTL effects in a locus including *Fam21* (C) and Il20ra (D) (red vertical stripes: proportion QTLs) and associated with variation in proportion of ILC1/ILC2 (Fam21 (C)), and ILC3/LTi (Il20ra (D)). (**E–G**) *Rpbj* cKO impact on ILC proportion and composition. (**E**) Left: Proportion of NKP46+ ILC3 (gated box) in the small intestine of wild-type *Rbpj*^+/+^*Rorc-* Cre (leftmost) and *Rbpj*^flox/flox^*Rorc*^Cre^ (second from left) mice. Right: Proportion of NKP46+ ILC3s (*y*-axis) in total CD45^+^ cells (second from right) and in ILCs (rightmost) in wildtype (black circles) and cKO (white circles) mice (*x*-axis). *n* = 6 individual mice in each category (dots). (**F**) Proportion CCR6-NKP46+ ILC3 in total ILC3 (*y*-axis) in wildtype (black circles) and cKO (white circles) mice (*x*-axis). *n* = 6 individual mice in each category (dots). (**G**) Left: Proportion of IL-17 producing ILC3 (gated box) in total ILC3 in the small intestine of wild-type *Rbp*j+/+*Rorc*^Cre^ (leftmost) and *Rbpj*^flox/flox^*Rorc*^Cre^ (second from left) mice. Right: Proportion of IL-17 producing ILC3 (*y*-axis) in total ILCs in wildtype (black circles) and cKO (white circles) mice (*x*-axis). *n* = 6 individual mice in each category (dots). Flow cytometry panels are representative of at least two independent experiments. Error bars: SEM. *P*-value: two-tailed *t* test.

**Fig. S3.**
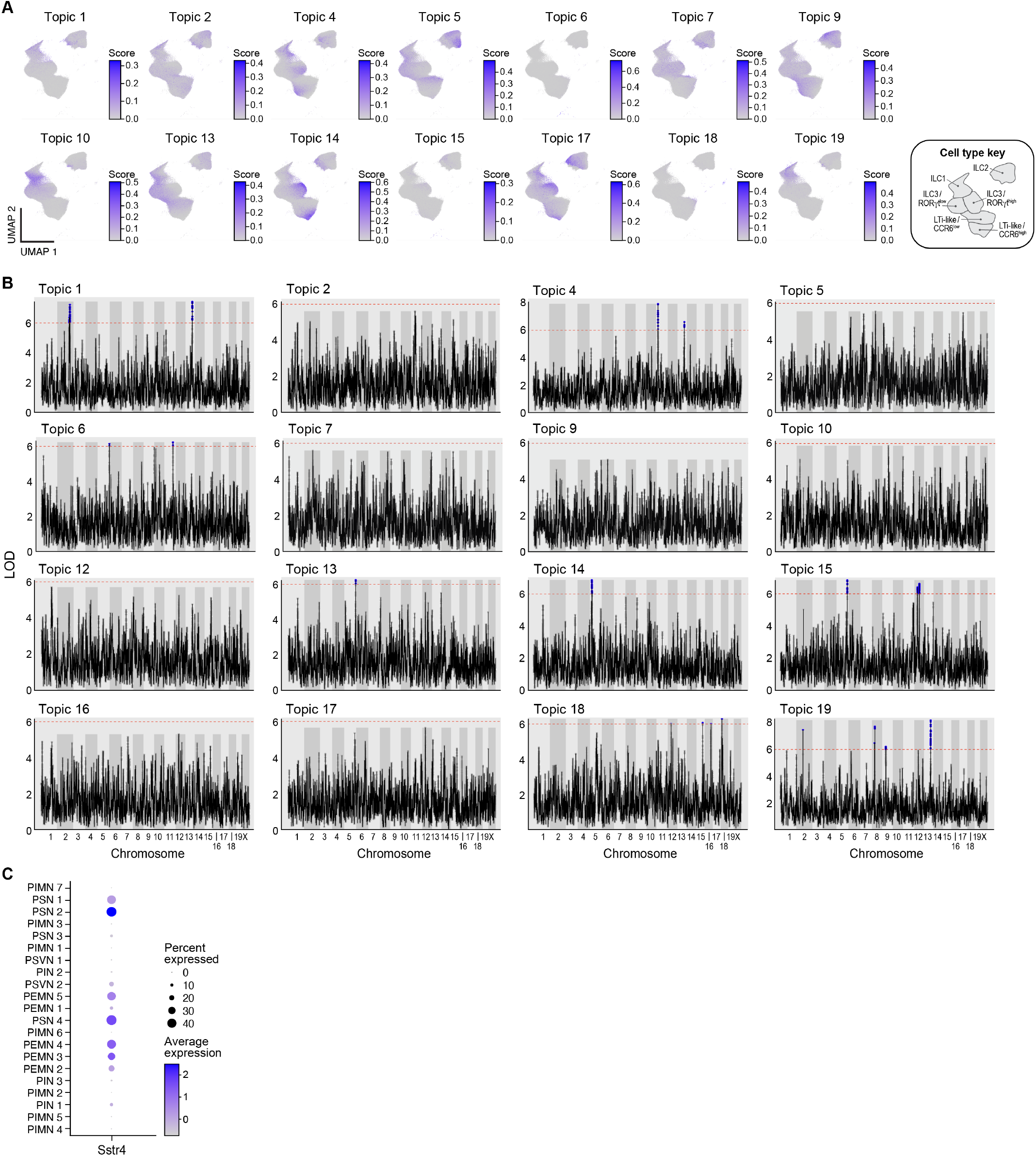
Programs and program QTLs. (**A**) Cross cell type programs. UMAP embedding of ILC profiles (as in Fig. 1C), colored by cell scores for each topic. Bottom right: cell type annotations. (**B**) Program QTLs. Significance (*y*-axis, –log_10_(*P*-value)) of association of each genomic locus (*x*-axis) with the cell score of a given topic (top left label). Dotted red line: genome-wide significance *p*-value = 0.05 (permutation-derived significance; Bonferroni corrected). Blue: significant associations. (**C**) *Sstr4* is expressed in mouse enteric neurons. Mean expression (dot color; log(TPM+1)) and proportion of expressing cells (dot size) of *Sstr4* across different subsets (rows) of mouse enteric neurons129.

**Fig. S4.**
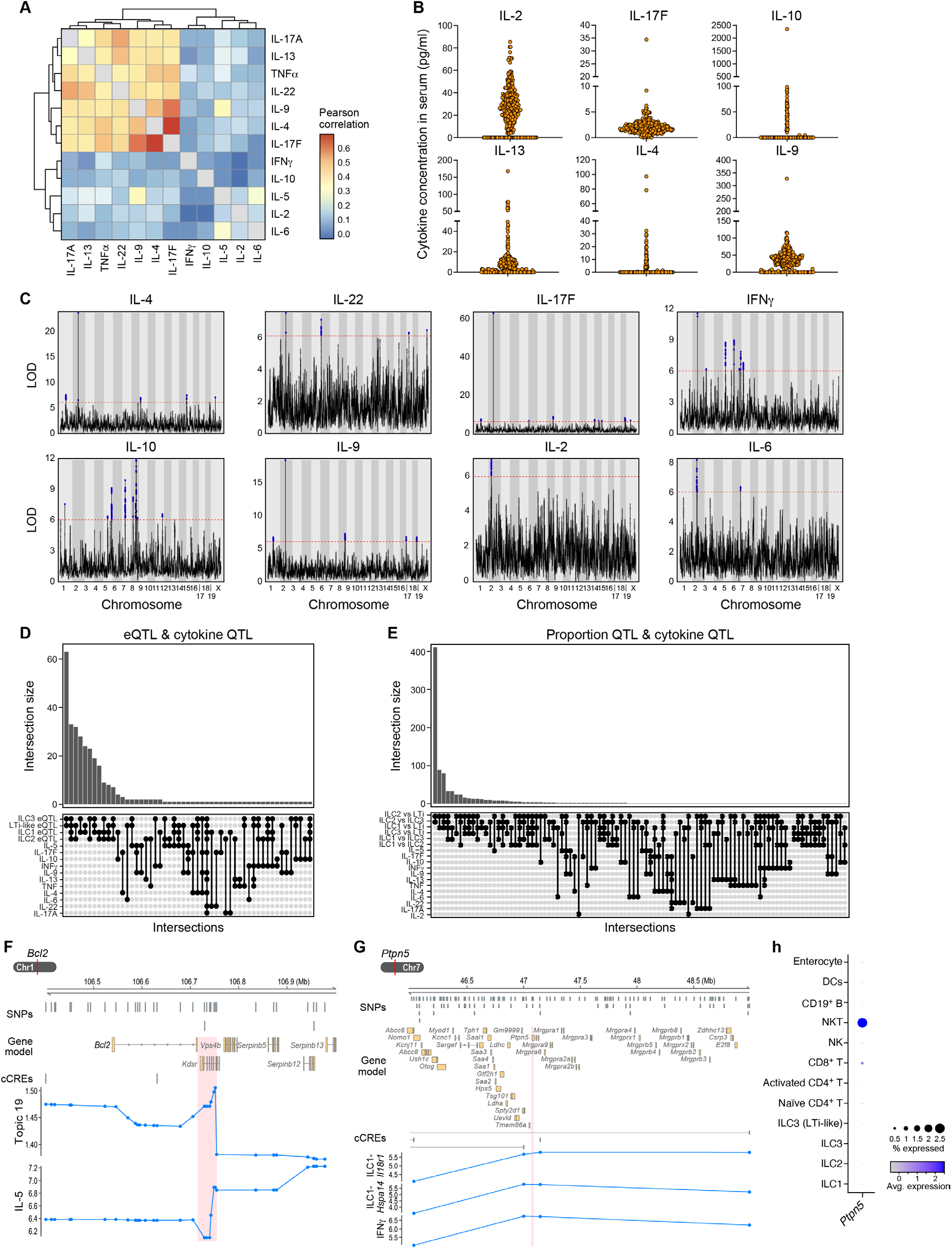
Serum cytokine QTLs. (**A**) Cytokine levels co-vary across mice. Correlation (color bar; Pearson’s *r*) in the level of different cytokines (rows, columns) across mice. (**B**) Variation in peripheral serum cytokine levels across 261 DO mice at homeostasis. Cytokine level (*y*-axis) for each cytokine (panel) in each mouse (dot). (**C**) Cytokine QTLs. LOD score (*y*-axis) for association of each genomic locus (*x*-axis) with a serum cytokine’s level (labeled on top). Dotted red line: *p*-value = 0.05 (Bonferroni corrected). Blue: significant associations. (**D–H**) Cytokine QTLs overlap with ILC eQTLs and proportion QTLs. (**D, E**) Number of loci (*y*-axis) shared between cytokine QTLs and specific categories of expression (**D**) or proportion (**E**) QTLs (*x*-axis, as marked on bottom). (**F**) Gene models (top), conserved regulatory elements (CRE) and LOD scores (*y*-axis) of QTL effects in a locus including *Bcl2, Kdsr* (red vertical stripe: cytokine QTL) and other genes and associated with variation in levels of cross-cell type topic 19 and serum IL-5. (**G**) Gene models (top), conserved regulatory elements (CRE) and LOD scores (*y*-axis) of QTL effects in a locus including *Ptpn5* and other genes (red vertical stripe: QTL) and associated with variation in ILC1 *Il18r1* and *Hspa14* and in serum IFNγ level. (**H**) Mean expression (dot color; log(TPM+1)) and proportion of expressing cells (dot size) of *Ptpn5* across different subsets (rows) of DO mice intestinal cells profiled by scRNA-seq.

**Fig. S5.**
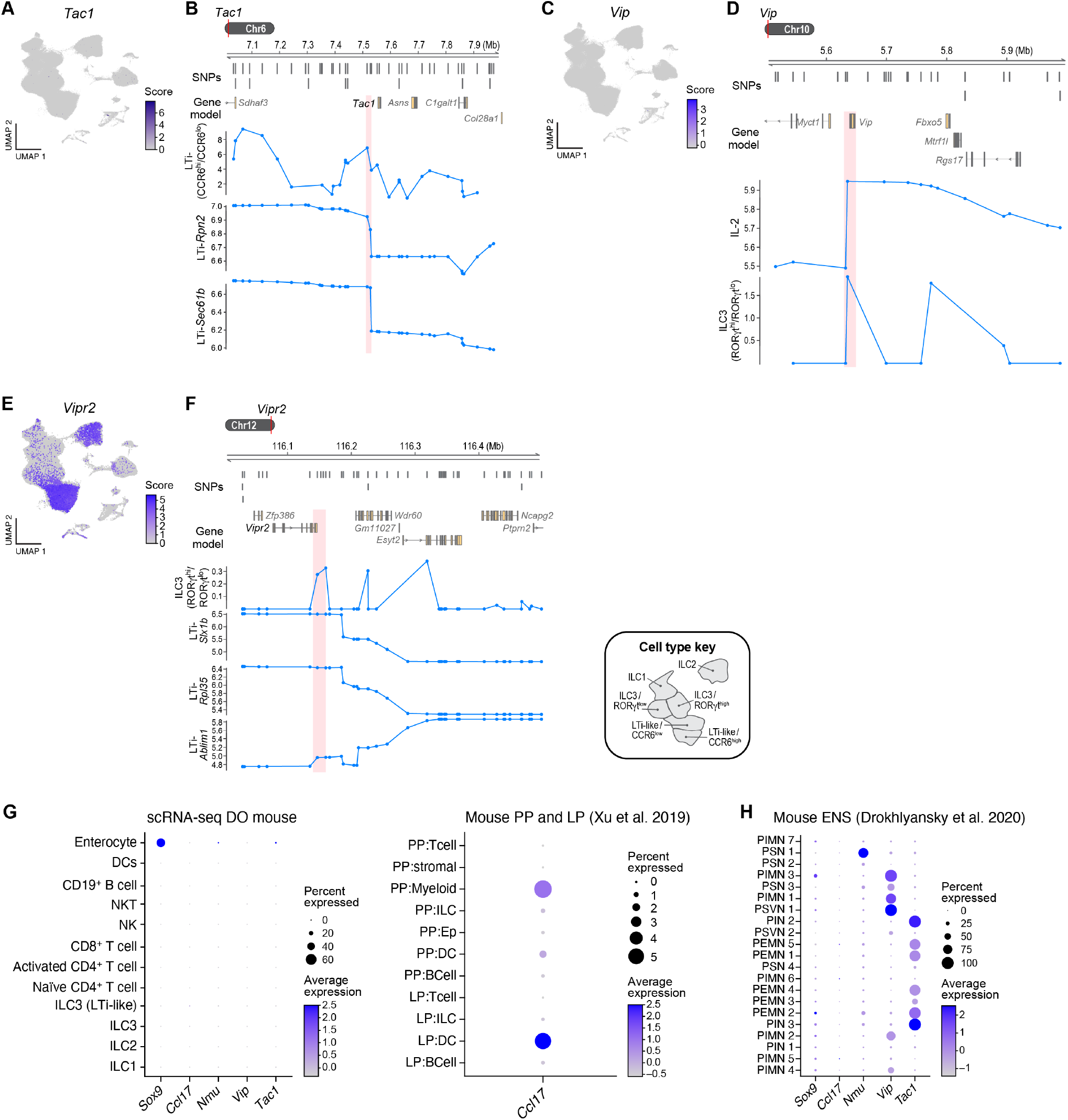
*Trans*-regulation cell type QTLs impacting ILC traits. (**A, B**) Neuron expressed *Tac1* associated with LTi-like cell proportion and *Sec61b* and *Rpn2* expression in ILC2s and ILC3s. (**A**) UMAP embedding of ILC profiles (as in Fig. 1C) colored by expression of *Tac1*. Inset: Cell type key. (**B**) Gene models (top) and LOD scores (*y*-axis) of QTL effects in a locus including *Tac1* and other genes (red vertical stripe: QTL) and associated with proportion of LTi (CCR6^high^/CCR6^low^) and *Sec61b* and *Rpn2* expression in LTis. (**C, D**) Neuron expressed *Vip* associated with proportion of ILC3 subset proportions and peripheral IL-2 level. (**C**) UMAP embedding of ILC profiles (as in Fig. 1C) colored by expression of *Vip*. (**D**) Gene models (top) and LOD scores (*y*-axis) of QTL effects in a locus including *Vip* and other genes (red vertical stripe: QTL) and associated with proportion of ILC3 (RORγt^high^/RORγt^low^) and peripheral IL-2 level. (**E, F**) ILC-expressed *Vipr2* associated with proportion of ILC3/LTi (RORγt^high^/RORγt^low^) and LTi gene expression. (**E**) UMAP embedding of ILC profiles (as in Fig. 1C) colored by expression of *Vipr2*. (**F**) Gene models (top) and LOD scores (*y*-axis) of QTL effects in a locus including *Vip* and other genes (red vertical stripe: QTL) and associated with proportion of ILC3 (RORγt^high^/RORγt^low^) and LTi expression of *Ablim1, Rpl35* and *Slx1b*. (**G, I**) Non-ILC expression of putative causal genes from DO QTLs. Mean expression (dot color; log(TPM+1)) and proportion of expressing cells (dot size) for key genes from highlighted QTLs (columns) across different cell subsets (rows) in DO mice (**G**, left), Peyer’s patches and the lamina propria10 (**G**, right), and the mouse enteric nervous system^129^ (**I**).

**Fig. S6.**
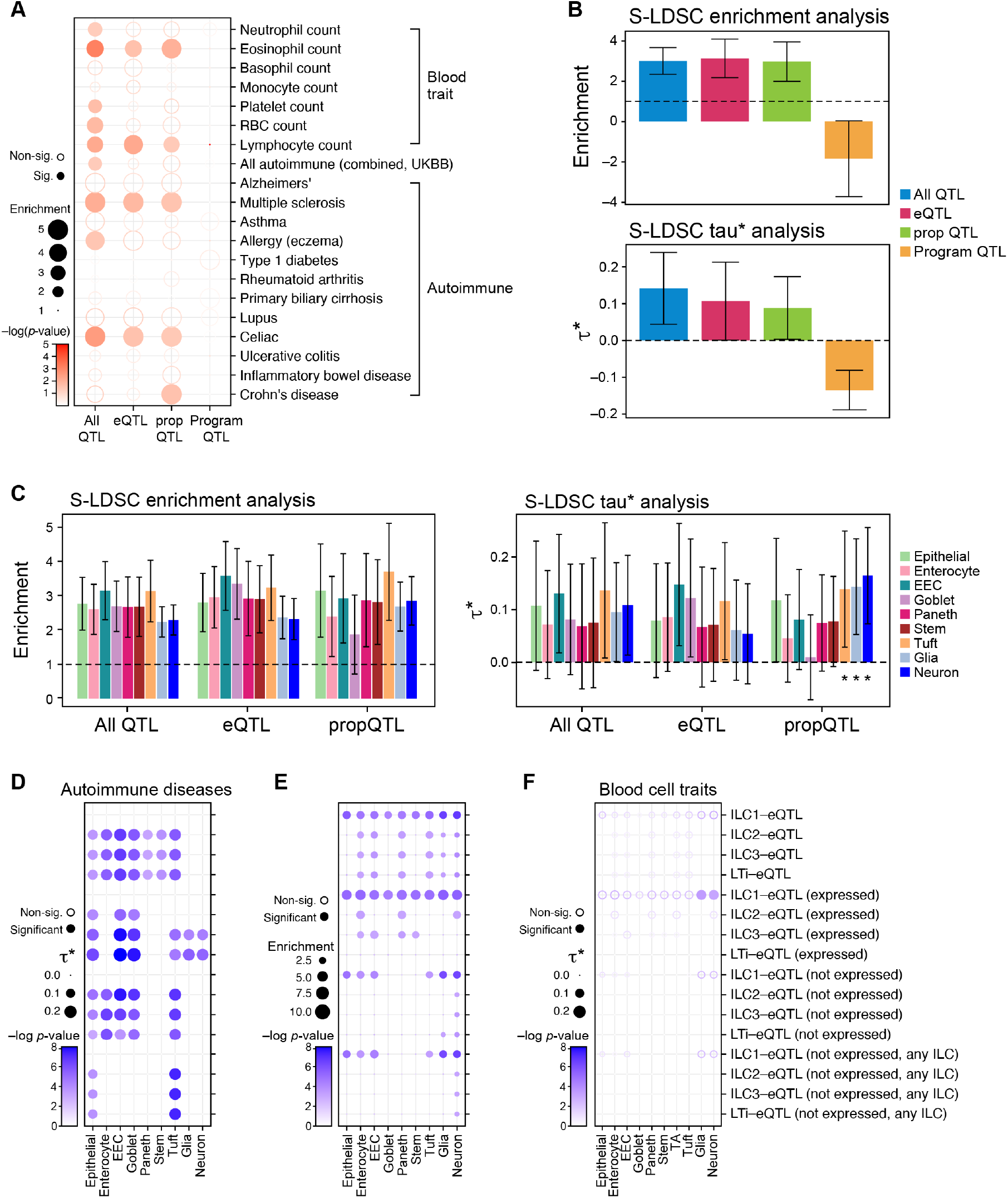
DO QTLs related to human disease risk heritability. (**A, B**) Heritability in human autoimmune and blood cell traits of human orthologs of genes in DO QTL categories. (**A**) sc-linker meta-analyzed heritability enrichments (top panel) and standardized effect sizes (bottom panel) across 7 relatively independent blood cell traits for human orthologs of mouse genes overlapping QTLs of different categories (*x*-axis). (**B**) Heatmap of sc-linker heritability enrichment signal of 7 blood cell traits and 12 autoimmune diseases for human orthologs of mouse genes overlapping QTLs from four categories (all types of QTLs, expression, proportion of ILC cell types and topic expression). Magnitude (enrichment, dot size) and significance (−log_10_(*P*), dot color). (**C–F**) Heritability in human autoimmune and blood cell traits of human orthologs of genes in DO QTL categories expressed in specific cell subsets. (**C**) sc-linker meta-analyzed heritability enrichments (left) and standardized effect sizes (right) across 12 relatively independent autoimmune diseases for human orthologs of mouse genes overlapping expression and proportion QTLs that are expressed in 9 other cell type classifications in mouse. (**D**) Meta-analyzed sc-linker conditional heritability signal represented by standardized effect sizes (τ*) across 12 autoimmune diseases for human orthologs of mouse genes overlapping eQTLs with 16 different characterizations of expression across 4 ILC cell sub-types (**Methods)** and expressed in 9 different mouse non-ILC cell types. Magnitude (τ*, dot size) and significance (−log_10_(*P*), dot color) of the standardized effect sizes are reported. (**E**) Meta-analyzed sc-linker heritability enrichments across 7 blood cell traits for human orthologs of mouse genes overlapping eQTLs with 16 different characterizations of expression across 4 ILC cell sub-types (Methods) and expressed in 9 different mouse non-ILC cell types. Magnitude (Enrichment, dot size) and significance (−log_10_(*P*), dot color) of the heritability enrichment are reported. (**F**) Meta-analyzed sc-linker conditional heritability signal represented by standardized effect sizes (τ*) across 7 blood cell traits for human orthologs of mouse genes overlapping eQTLs with 16 different characterizations of expression across 4 ILC cell sub-types (**Methods)** and expressed in 9 different mouse non-ILC cell types. Magnitude (enrichment, dot size) and significance (−log_10_(*P*), dot color) of the heritability enrichment are reported. All heritability results are conditional on the 87 baseline-LD (v2.1) annotations. For sc-linker analysis, enhancer-gene links were aggregated across all tissues.

## Notes

### Competing Interest Statement

The authors declare competing financial interests: A.R. is a cofounder and equity holder of Celsius Therapeutics, an equity holder in Immunitas, and was an SAB member of ThermoFisher Scientific, Syros Pharmaceuticals, Neogene Therapeutics and Asimov until July 31, 2020. From August 1, 2020, A.R. is an employee of Genentech. R.J.X. is a cofounder of Celsius Therapeutics, Jnana Therapeutics, and a member of the scientific advisory board of Magnet Bio medicine, Nestle and Moonlake Immunotherapeutics.

